# On the neural substrates of mind wandering and dynamic thought: A drug and brain stimulation study

**DOI:** 10.1101/2024.11.02.620526

**Authors:** Tara Rasmussen, Paul E. Dux, Hannah Filmer

## Abstract

The impact of mind wandering on our daily lives ranges from diminishing productivity, to facilitating creativity and problem solving. There is evidence that distinct internal thought types can be modulated by transcranial direct current stimulation (tDCS), although little is known about optimal stimulation parameters or the mechanisms behind such effects. In addition, recent findings suggest changes in dopamine availability may alter the effect tDCS has on neural and behavioural outcomes. Dopaminergic functioning has also been implicated in executive processes anticorrelated with mind wandering such as attention and working memory, however the neurochemical mechanisms involved in internal thoughts are largely unknown. Here, we investigated the role of dopamine, and tDCS, on internal thought processes. Specifically, using an attentional control task, we tested whether dopamine availability (levodopa or placebo) mediated the effects of online high definition tDCS (HD-tDCS; 2mA, or sham). There was no evidence for our hypothesised effect of left prefrontal cortex HD-tDCS reducing task unrelated thought, nor freely moving thought. This failure to replicate previous HD-tDCS findings emphasises the importance of employing robust methodological practices within this field to improve confidence in the findings. However, we did find that levodopa reduced freely moving thought, relative to placebo. We also found preliminary evidence that dopamine availability may moderate the relationship between stimulation and behavioural variability performance during periods of task unrelated thought. Overall, these findings suggest that stimulation does not affect dynamic internal thought, however there is initial evidence for the potential effectiveness of targeting the dopaminergic system to reduce spontaneous internal thoughts and improve behavioural performance.

---

Cognitive control and mind wandering represent the “yin and yang” of executive function. Mind wandering – the direction of thoughts towards self-generated, internally orientated representations – is a complex phenomenon, and this heterogeneity has been recently characterised in the dynamic framework (Martel et al., 2019). This hypothesis suggests there are three dynamic thought types – deliberately constrained thoughts, automatically constrained thoughts and freely moving thoughts (Kam et al., 2021; Martel et al., 2019; Seli, Kane, et al., 2018). Currently, little is known on the neural substrates underlying these distinct thought types, however, neuroimaging has indicated that similar neural networks underlie mind wandering and cognitive control operations, particularly those implicated in maintaining focus on goal directed representations (Christoff et al., 2016; Fox et al., 2015; Groot et al., 2021). Nevertheless, the underlying causal neural mechanisms which drive a shift from task focussed towards internally oriented thoughts remain poorly understood.

Non-invasive brain stimulation approaches – such as transcranial direct current stimulation (tDCS) – can be applied to understand the causal neural substrates associated with mind wandering and attentional control. tDCS works by passing a weak electrical current (typically between 0.5mA and 4mA) between electrodes which are placed on the scalp (Filmer et al., 2014, 2020). High definition (HD) tDCS (see Figure 1), uses small electrodes, typically arranged in a 4 x 1 ring montage, to pass the current from the central anodal electrode to the four surrounding reference cathodes (Datta et al., 2009; Villamar et al., 2013). Consistent with imaging research, tDCS studies have causally implicated the prefrontal cortex (PFC) in mind wandering (Axelrod et al., 2018; Boayue et al., 2021; Filmer et al., 2019). But an early research finding that 1mA anodal tDCS applied to the left PFC increased mind wandering (Axelrod et al., 2015, 2018) has failed to replicate in a high-powered replication study which found strong evidence against a stimulation effect (Boayue et al., 2020). Furthermore, studies applying 2mA HD-tDCS to the PFC have also shown both support for (Boayue et al., 2021) and against (Alexandersen et al., 2022) modulations to mind wandering. Regarding the dynamic framework, we recently found preliminary evidence, in a high-powered registered report, that freely moving thoughts were reduced by 2mA HD-tDCS stimulation being applied to the left PFC and deliberately constrained thoughts were reduced by stimulation to the right inferior parietal lobule (IPL; (Rasmussen et al., 2024). These findings suggest dynamic thought types can be modulated by HD-tDCS and have potentially distinct neural substrates; however, the mechanisms behind such modulations remain unclear.

**Figure 1.**
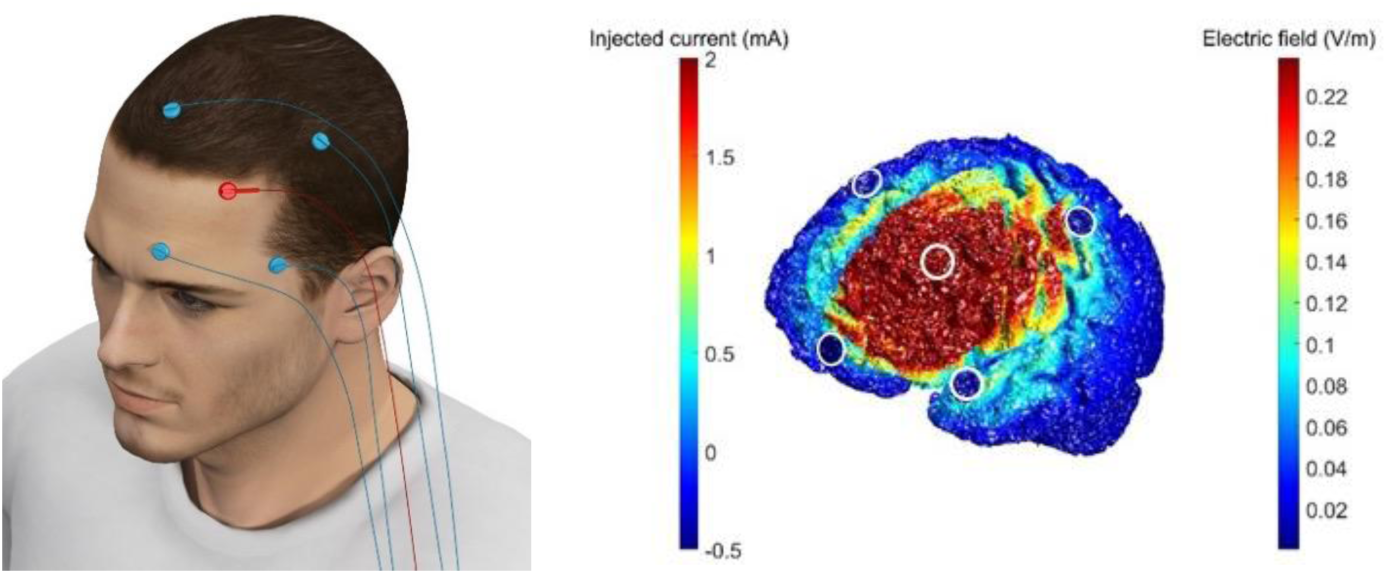
HD-tDCS montage and current modelling. Display of electrode placement over the left PFC, with the anode at F3 and cathodes placed at F7, C3, Fz and Fp1 (left image) and current modelling for this 4 x 1 ring HD-tDCS montage (right image).

Brain stimulation has been found to directly affect the excitability of various neurochemical mechanisms, including inducing changes in the concentration of dopamine neurotransmitters in cortical and subcortical regions (Bunai et al., 2021; Fonteneau et al., 2018; Fukai et al., 2019; Meyer et al., 2019). The importance of the dopaminergic system in cognitive control has been consistently highlighted, whereby dopamine manipulations have been shown to exert a dosage-dependent, inverted U-shaped effect, on attention and working memory processes (Cools, 2016; Cools & D’Esposito, 2011; D’Ardenne et al., 2012). Furthermore, there is evidence to suggest effects of PFC brain stimulation on cognitive control processes are related to changes in dopamine concentrations. For example, research has found tDCS induced improvements in the accuracy component of an executive functioning task were correlated with an increase of dopamine released in the right ventral striatum, which is linked to PFC through the meso-cortical-limbic system (Bunai et al., 2021; Fukai et al., 2019). There is also evidence to suggest this dopamine-tDCS interaction may directly influence behavioural outcomes (Borwick et al., 2020; Leow, Jiang, et al., 2023; Leow, Marcos, et al., 2023). This research highlights the influence of dopamine on brain stimulation outcomes, however, no research to date has investigated the causal role of dopamine in internal thought processes.

## The present study

The current study employed an anodal HD-tDCS protocol, applied to the left PFC, in conjunction with a levodopa manipulation, designed to increase dopamine availability, to explore the interaction between dopamine and stimulation effects on mind wandering, during an attention control task. Table 1 provides a full summary of the study design.

**Table 1.** The demographic information which was used to balance the six groups of participants. This includes age, sex, time spent on video games, time spent playing musical instruments and hours of sleep from the night before.

|  | Active stimulation<br>with levodopa |  | Active stimulation<br>with placebo |  | Sham stimulation<br>with levodopa |  | Sham stimulation<br>with placebo |  |
| --- | --- | --- | --- | --- | --- | --- | --- | --- |
|  | *M* (SD) | *n* | *M* (SD) | *n* | *M* (SD) | *n* | *M* (SD) | *n* |
| **Age** | 21.51<br>(4.36) | 59 | 21.74<br>(3.68) | 58 | 21.49<br>(2.78) | 57 | 21.60<br>(4.42) | 57 |
| **Sex (*n*, %**<br>**female)** | 44<br>(74.58%) | 59 | 43<br>(74.14%) | 58 | 43<br>(75.44%) | 57 | 44<br>(77.19%) | 57 |
| **Time spent on**<br>**video games** | 2.20<br>(1.48) | 59 | 2.28<br>(1.48) | 58 | 2.37<br>(1.73) | 57 | 2.25<br>(1.50) | 57 |
| **Time spent on**<br>**musical**<br>**instruments** | 1.34<br>(0.80) | 59 | 1.33<br>(0.69) | 58 | 1.37<br>(0.79) | 57 | 1.37<br>(0.88) | 57 |
| **Hours of sleep** | 7.09<br>(1.31) | 59 | 7.12<br>(1.10) | 58 | 7.00<br>(1.26) | 57 | 7.11<br>(1.19) | 57 |

This research first aimed to replicate the effect of PFC stimulation on freely moving found by Rasmussen et al. (2024), whereby we hypothesised that 2mA anodal HD-tDCS to the left PFC would reduce freely moving thought, relative to the sham group, across participants in the placebo drug condition (H_1a_). While there have been contrasting findings on the effect of tDCS on task unrelated thought, there is evidence to suggest that stimulation can also reduce these thoughts (Boayue et al., 2021), thus we hypothesised that 2mA anodal HD-tDCS would reduce task unrelated thought, relative to the sham group, across the placebo drug groups (H_1b_). To understand the neurochemical mechanisms underlying mind wandering and the dynamic thought types, this research also aimed to investigate whether the changes in mind wandering and dynamic thought, while completing a cognitively demanding task, were being driven by changes in dopamine availability. We hypothesised that there would be an effect of increasing dopamine availability via levodopa, compared to the placebo group, on freely moving thought (a) and task unrelated thought (b), across the sham stimulation conditions (H_2_). Finally, because there is evidence that levodopa may mediate the effects of tDCS on performance outcomes (Leow, Jiang, et al., 2023; Leow, Marcos, et al., 2023) we also aimed to investigate how the interaction between tDCS and dopamine affected internal thought types in the PFC. Thus, we predicted there would be a difference in the effect of 2mA HD-tDCS in combination with levodopa on freely moving thought (a) and task unrelated thought (b), relative to the active placebo groups (H_3_).

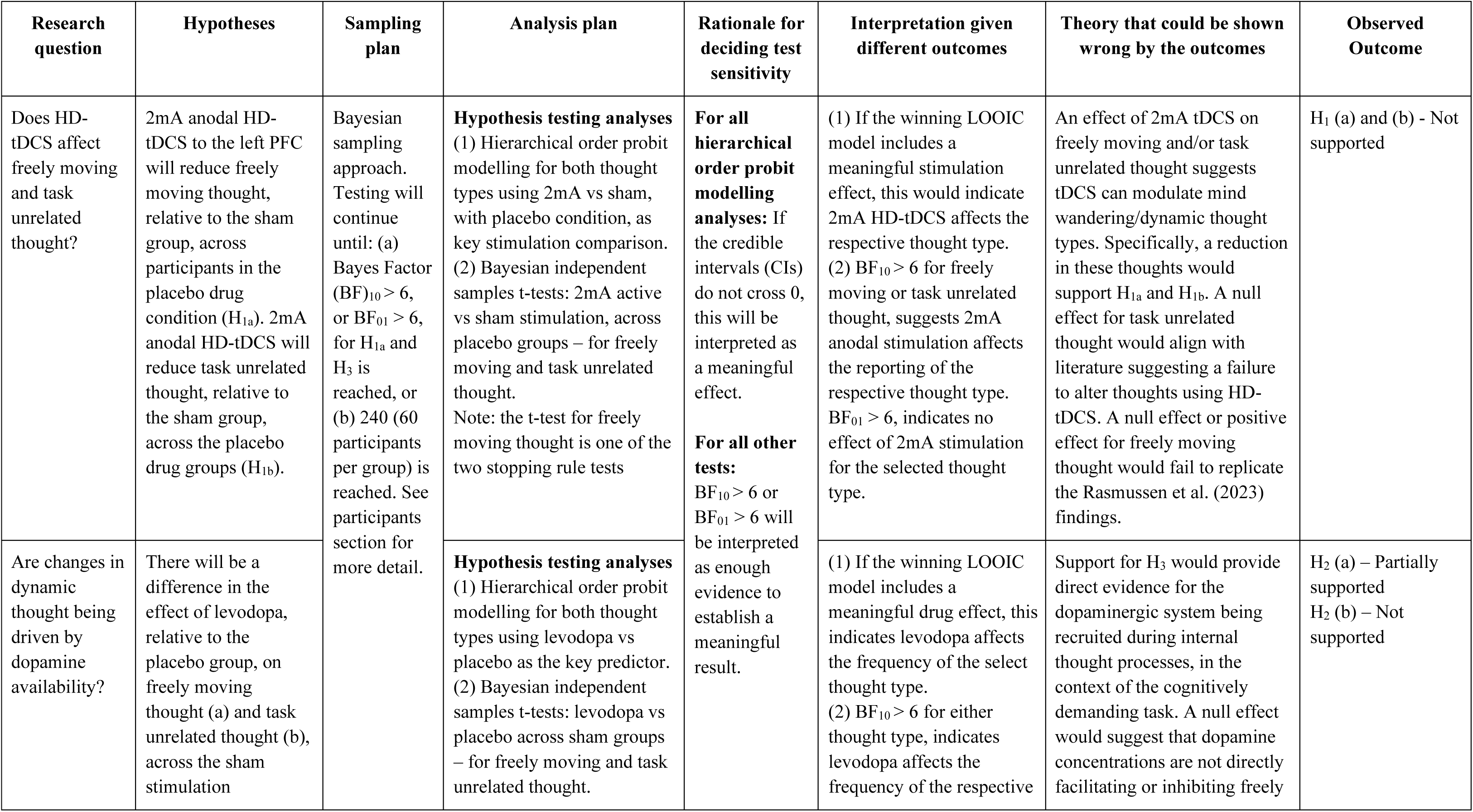
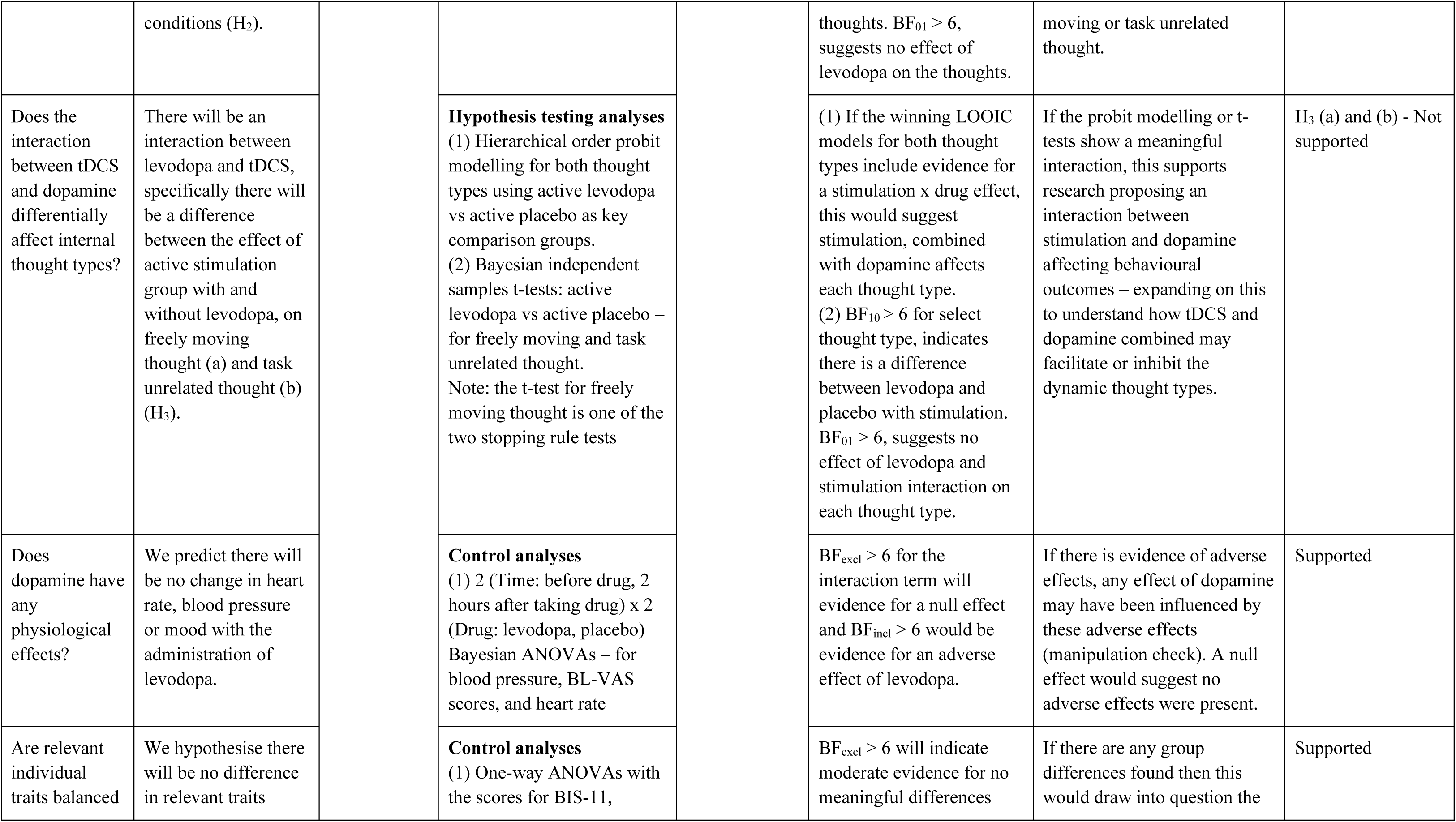
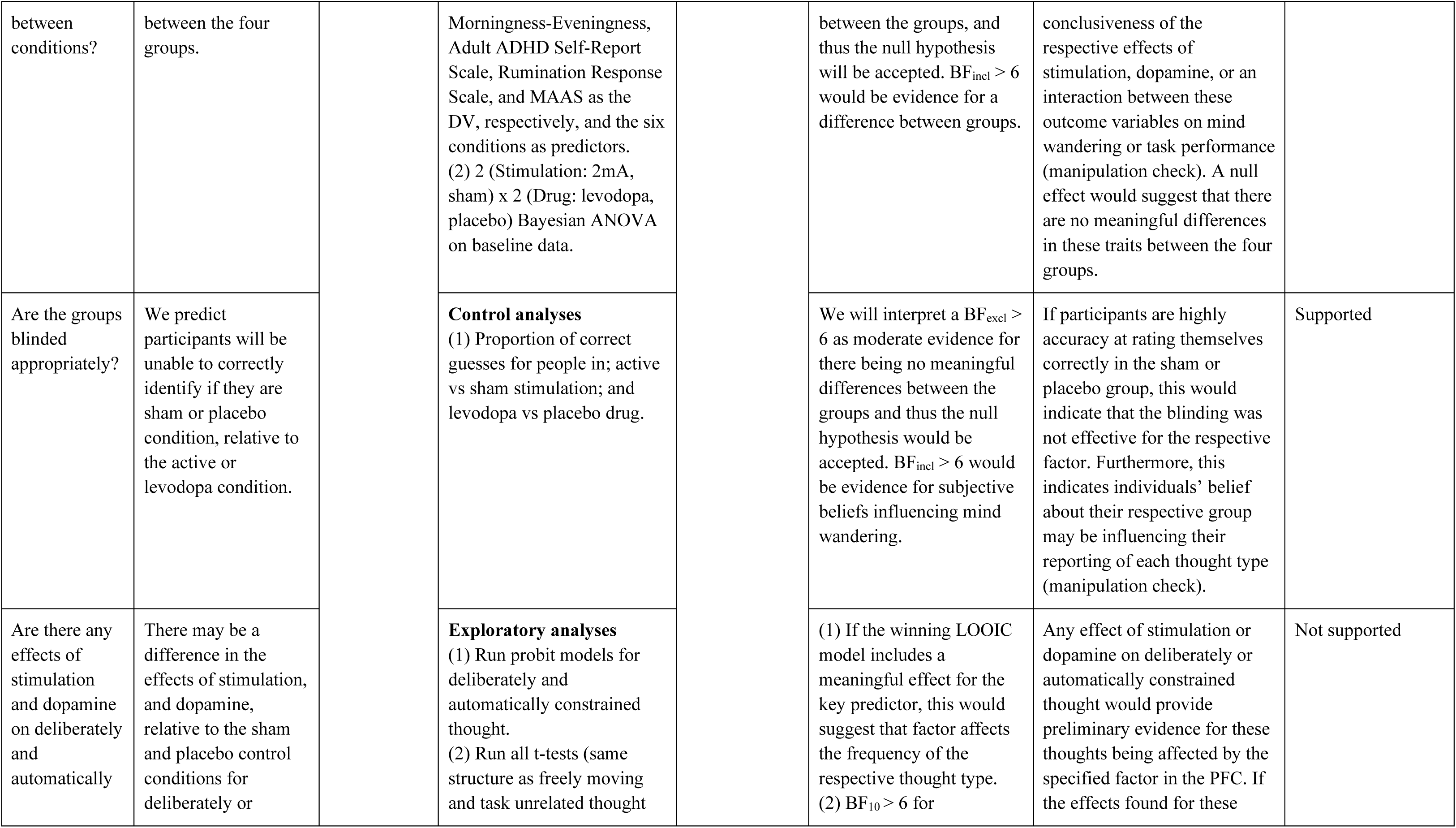
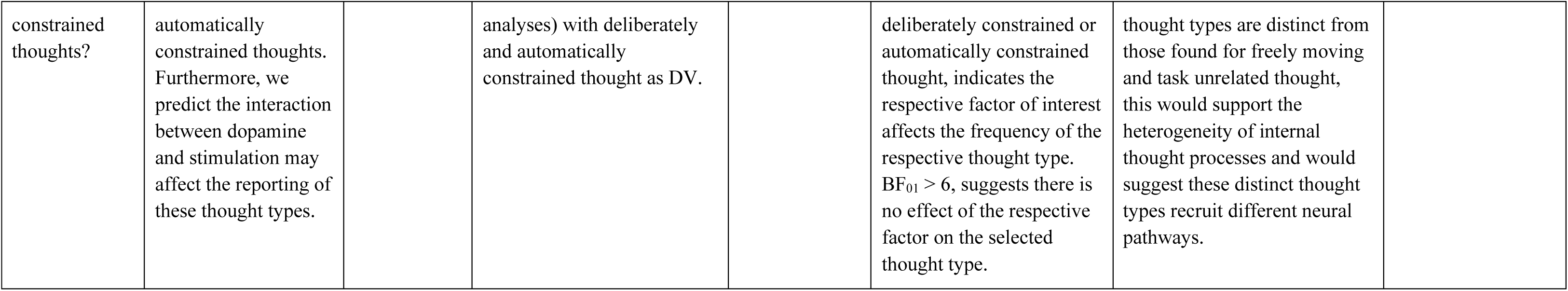

| Research question | Hypotheses | Sampling plan | Analysis plan | Rationale for deciding test sensitivity | Interpretation given different outcomes | Theory that could be shown wrong by the outcomes | Observed Outcome |
| --- | --- | --- | --- | --- | --- | --- | --- |
| Does HD-tDCS affect freely moving and task unrelated thought? | 2mA anodal HD-tDCS to the left PFC will reduce freely moving thought, relative to the sham group, across participants in the placebo drug condition ($H_{1a}$). 2mA anodal HD-tDCS will reduce task unrelated thought, relative to the sham group, across the placebo drug groups ($H_{1b}$). | Bayesian sampling approach. Testing will continue until: (a) Bayes Factor $(BF)_{10} > 6$, or $BF_{01} > 6$, for $H_{1a}$ and $H_3$ is reached, or (b) 240 (60 participants per group) is reached. See participants section for more detail. | **Hypothesis testing analyses**<br>(1) Hierarchical order probit modelling for both thought types using 2mA vs sham, with placebo condition, as key stimulation comparison.<br>(2) Bayesian independent samples t-tests: 2mA active vs sham stimulation, across placebo groups – for freely moving and task unrelated thought.<br>Note: the t-test for freely moving thought is one of the two stopping rule tests | **For all hierarchical order probit modelling analyses:** If the credible intervals (CIs) do not cross 0, this will be interpreted as a meaningful effect.<br>**For all other tests:** $BF_{10} > 6$ or $BF_{01} > 6$ will be interpreted as enough evidence to establish a meaningful result. | (1) If the winning LOOIC model includes a meaningful stimulation effect, this would indicate 2mA HD-tDCS affects the respective thought type.<br>(2) $BF_{10} > 6$ for freely moving or task unrelated thought, suggests 2mA anodal stimulation affects the reporting of the respective thought type.<br>$BF_{01} > 6$, indicates no effect of 2mA stimulation for the selected thought type. | An effect of 2mA tDCS on freely moving and/or task unrelated thought suggests tDCS can modulate mind wandering/dynamic thought types. Specifically, a reduction in these thoughts would support $H_{1a}$ and $H_{1b}$. A null effect for task unrelated thought would align with literature suggesting a failure to alter thoughts using HD-tDCS. A null effect or positive effect for freely moving thought would fail to replicate the Rasmussen et al. (2023) findings. | $H_1$ (a) and (b) - Not supported |
| Are changes in dynamic thought being driven by dopamine availability? | There will be a difference in the effect of levodopa, relative to the placebo group, on freely moving thought (a) and task unrelated thought (b), across the sham stimulation | | **Hypothesis testing analyses**<br>(1) Hierarchical order probit modelling for both thought types using levodopa vs placebo as the key predictor.<br>(2) Bayesian independent samples t-tests: levodopa vs placebo across sham groups – for freely moving and task unrelated thought. | | (1) If the winning LOOIC model includes a meaningful drug effect, this indicates levodopa affects the frequency of the select thought type.<br>(2) $BF_{10} > 6$ for either thought type, indicates levodopa affects the frequency of the respective | Support for $H_3$ would provide direct evidence for the dopaminergic system being recruited during internal thought processes, in the context of the cognitively demanding task. A null effect would suggest that dopamine concentrations are not directly facilitating or inhibiting freely | $H_2$ (a) – Partially supported<br>$H_2$ (b) – Not supported |
| | conditions (H<sub>2</sub>). | | | | thoughts. $BF_{01} > 6$, suggests no effect of levodopa on the thoughts. | moving or task unrelated thought. | |
| Does the interaction between tDCS and dopamine differentially affect internal thought types? | There will be an interaction between levodopa and tDCS, specifically there will be a difference between the effect of active stimulation group with and without levodopa, on freely moving thought (a) and task unrelated thought (b) (H<sub>3</sub>). | | **Hypothesis testing analyses**<br>(1) Hierarchical order probit modelling for both thought types using active levodopa vs active placebo as key comparison groups.<br>(2) Bayesian independent samples t-tests: active levodopa vs active placebo – for freely moving and task unrelated thought.<br>Note: the t-test for freely moving thought is one of the two stopping rule tests | | (1) If the winning LOOIC models for both thought types include evidence for a stimulation x drug effect, this would suggest stimulation, combined with dopamine affects each thought type.<br>(2) $BF_{10} > 6$ for select thought type, indicates there is a difference between levodopa and placebo with stimulation. $BF_{01} > 6$, suggests no effect of levodopa and stimulation interaction on each thought type. | If the probit modelling or t-tests show a meaningful interaction, this supports research proposing an interaction between stimulation and dopamine affecting behavioural outcomes – expanding on this to understand how tDCS and dopamine combined may facilitate or inhibit the dynamic thought types. | H<sub>3</sub> (a) and (b) - Not supported |
| Does dopamine have any physiological effects? | We predict there will be no change in heart rate, blood pressure or mood with the administration of levodopa. | | **Control analyses**<br>(1) 2 (Time: before drug, 2 hours after taking drug) x 2 (Drug: levodopa, placebo) Bayesian ANOVAs – for blood pressure, BL-VAS scores, and heart rate | | $BF_{excl} > 6$ for the interaction term will evidence for a null effect and $BF_{incl} > 6$ would be evidence for an adverse effect of levodopa. | If there is evidence of adverse effects, any effect of dopamine may have been influenced by these adverse effects (manipulation check). A null effect would suggest no adverse effects were present. | Supported |
| Are relevant individual traits balanced | We hypothesise there will be no difference in relevant traits | | **Control analyses**<br>(1) One-way ANOVAs with the scores for BIS-11, | | $BF_{excl} > 6$ will indicate moderate evidence for no meaningful differences | If there are any group differences found then this would draw into question the | Supported |
| between conditions? | between the four groups. | | Morningness-Eveningness, Adult ADHD Self-Report Scale, Rumination Response Scale, and MAAS as the DV, respectively, and the six conditions as predictors.<br>(2) 2 (Stimulation: 2mA, sham) x 2 (Drug: levodopa, placebo) Bayesian ANOVA on baseline data. | | between the groups, and thus the null hypothesis will be accepted. $BF_{incl} > 6$ would be evidence for a difference between groups. | conclusiveness of the respective effects of stimulation, dopamine, or an interaction between these outcome variables on mind wandering or task performance (manipulation check). A null effect would suggest that there are no meaningful differences in these traits between the four groups. | |
| Are the groups blinded appropriately? | We predict participants will be unable to correctly identify if they are sham or placebo condition, relative to the active or levodopa condition. | | **Control analyses**<br>(1) Proportion of correct guesses for people in; active vs sham stimulation; and levodopa vs placebo drug. | | We will interpret a $BF_{excl} > 6$ as moderate evidence for there being no meaningful differences between the groups and thus the null hypothesis would be accepted. $BF_{incl} > 6$ would be evidence for subjective beliefs influencing mind wandering. | If participants are highly accurate at rating themselves correctly in the sham or placebo group, this would indicate that the blinding was not effective for the respective factor. Furthermore, this indicates individuals' belief about their respective group may be influencing their reporting of each thought type (manipulation check). | Supported |
| Are there any effects of stimulation and dopamine on deliberately and automatically | There may be a difference in the effects of stimulation, and dopamine, relative to the sham and placebo control conditions for deliberately or | | **Exploratory analyses**<br>(1) Run probit models for deliberately and automatically constrained thought.<br>(2) Run all t-tests (same structure as freely moving and task unrelated thought | | (1) If the winning LOOIC model includes a meaningful effect for the key predictor, this would suggest that factor affects the frequency of the respective thought type.<br>(2) $BF_{10} > 6$ for | Any effect of stimulation or dopamine on deliberately or automatically constrained thought would provide preliminary evidence for these thoughts being affected by the specified factor in the PFC. If the effects found for these | Not supported |
| constrained thoughts? | automatically constrained thoughts. Furthermore, we predict the interaction between dopamine and stimulation may affect the reporting of these thought types. | | analyses) with deliberately and automatically constrained thought as DV. | | deliberately constrained or automatically constrained thought, indicates the respective factor of interest affects the frequency of the respective thought type. $BF_{01} > 6$, suggests there is no effect of the respective factor on the selected thought type. | thought types are distinct from those found for freely moving and task unrelated thought, this would support the heterogeneity of internal thought processes and would suggest these distinct thought types recruit different neural pathways. | |

## Methodology

### Ethics approval

This research was approved by The University of Queensland Human Research Ethics Committee (Clearance ID: 2009000335). Participants were recruited through the paid participant pool, the School of Psychology first-year volunteer pool at The University of Queensland, and flyers. Compensation was either $20 per hour or course credits. All participants were required to provide informed consent to be eligible to participate in the study.

### Design

Each participant completed a single session, which consisted of either 2mA active or sham HD-tDCS, in conjunction with a dopaminergic manipulation (levodopa or placebo drug). The stimulation was delivered in conjunction with the Finger-Tapping Random-Sequence Generation Task (FT-RSGT), which is designed for participants to generate random sequences in time to a metronome tone. This study employed a between subjects’ design, whereby participants were pseudo-randomly allocated to one group according to the following variables: drug (levodopa, placebo) x stimulation condition (anodal HD-tDCS, sham HD-tDCS). Thus, the conditions were: (1) sham HD-tDCS and placebo; (2) sham HD-tDCS and levodopa; (3) 2mA anodal HD-tDCS and placebo; (4) 2mA anodal HD-tDCS and levodopa. A between-subjects approach was most appropriate for this study as it helped preserve the integrity of the stimulation and dopamine blinding. Further, it reduced the likelihood of practice effects in the task or any inter-session changes, which are associated with within-subjects designs. The session began with participants completing a demographic questionnaire which was used to pseudo-randomly allocate participants into demographically balanced groups. This was conducted in a double-blinded manner via a MATLAB allocation script. This method was also designed to reduce the likelihood of group-related confounds in the between-groups design. They then received training on the FT-RSGT task and the four thought probes that were presented throughout, before completing a 10-minute baseline block on the task (see Figure 2). The levodopa or placebo drug was then crushed and mixed with orange juice for participants to consume and administered by an alternative experimenter, while the HD-tDCS electrodes were set up. This was to ensure the drug manipulation was blinded for both the participants and experimenter. During this period, participants also completed some trait-based questionnaires. After 40 minutes passed since the drug administration, participants then completed a 30-minute stimulation block, consisting of the FT-RSGT in conjunction with online active or sham stimulation. Once they completed the task, an end of session questionnaire was administered, and participants were debriefed on their participation in the study. The full Stage 1 registered report manuscript can also be accessed on the OSF (https://osf.io/an57y/).

**Figure 2.**
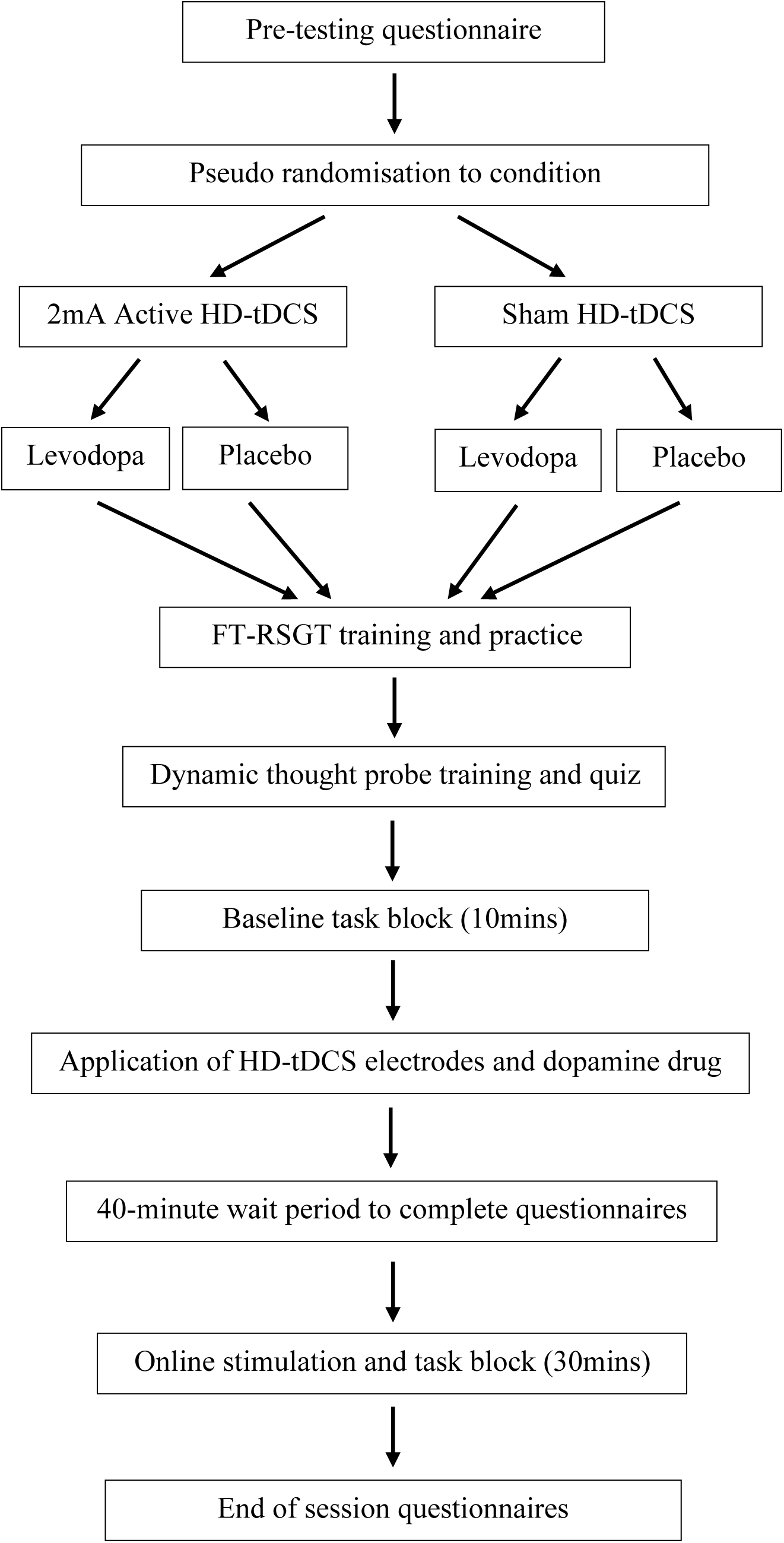
Experimental procedure. The sessions begun with a pre-screening and pseudo random allocation to one of four stimulation and drug groups, which was followed by training and a 10-minute baseline block. The electrodes were then applied, in conjunction with the dopamine drug manipulation. After a 40-minute wait period, participants then completed a 30-minute online stimulation block and finally an end of session questionnaire and debriefing on the experiment.

### Participants

#### Bayesian sampling plan

There was no pre-determined sample size for this study, as it employed a Bayesian sampling approach. The study recruited a total of 252 participants; however, 12 participants were removed from the sample prior to any analysis. Specifically, and in line with our exclusion criteria: five participants were removed due to issues relating to the stimulation (e.g. the participant withdrew due to the sensations they experienced); four participants were removed as they did not complete the task correctly; and despite pre-screening, three participants were found to be ineligible at the start of their session. This resulted in a dataset consisting of 240 participants. After the testing was completed, a further 9 participants were removed, according to our post-study data exclusions outlined in the Analysis Overview below. This resulted in a final sample of 231 participants.

The Bayesian sampling approach stated participants would continue to be recruited until a Bayes Factor (BF)_10_ > 6 or BF_01_ > 6 was reached for the selected hypothesis tests (see above). However, these conditions were not met, thus recruitment continued until the maximum sample size of 240 complete datasets (60 participants per group; the maximum number dictated by resource constraints) was reached. This was larger than the sample size which has been used previously to find meaningful results (Rasmussen et al., 2024) and we believed inconclusive results in the chosen tests at this sample size would still offer an important contribution to the literature. The stopping rule was first checked after 15 participants in each group were tested, and for every 5 participants per group thereafter. The first selected hypothesis test was that freely moving thought would be reduced with active HD-tDCS, relative to sham group, for the placebo conditions. Specifically, we ran a Bayesian independent samples t-test which compared the active group to the sham stimulation group, for participants in the placebo condition. This was run on the stimulation block data alone, with average freely moving thought responses as the dependent variable. This test was selected to replicate the effect found by Rasmussen et al. (2024), whereby 2mA HD-tDCS to the left PFC decreased freely moving thought relative to sham. The additional selected hypothesis test was that there would be an interaction between levodopa and tDCS, specifically there would be a difference between the effect of active stimulation groups with and without levodopa, on freely moving thought. This was assessed using a Bayesian independent samples t-test which compared the active levodopa group to the active placebo group on the stimulation block data alone, with the average freely moving thought responses as the dependent variable. This additional test was selected as there is evidence levodopa may mediate the effects of tDCS on performance outcomes (Leow, Jiang, et al., 2023; Leow, Marcos, et al., 2023), thus we believed finding an interaction between these factors would provide a meaningful contribution to the field.

#### Exclusion criteria

Participants (aged 18-40) were required to be right handed and to meet the following criteria, in order to be included in the study: (1) English as their primary language; (2) No current use of psychiatric medication(s); (3) No current or previous psychiatric/neurological condition(s); (4) No current use of psychotropic drugs or blood pressure medication; (5) Not currently taking part in other tDCS studies and they had to meet the tDCS safety screening questionnaire criteria (see supplementary materials; e.g. no implanted medical device or metal in the head). Participants were also excluded and replaced in the data collection phase if they could not understand the thought probes after training and additional clarification from the experimenter. After completing the screening and training on the task, participants were pseudo-randomly allocated to demographically balanced groups using a MATLAB script, which allowed for the double-blinding of each subject’s condition. This script accounted for participants age, sex, time spent playing video games and musical instruments, hours of sleep from the previous night and whether their session was before or after 12:00pm (i.e., AM or PM; see Table 1 for demographic information).

Participants were also excluded from the study and replaced during the testing phase if their responses to the end of session questionnaire suggested that the participant did not understand how to correctly generate random number sequences. Specifically, if participants cited that they used the same pattern throughout which repeated more than twice at a time (e.g. *z,z,z,m,m,z,z,z,m,m*) or if they stated that they only alternated from one key to the other in the same order throughout, with one or more taps at a time on each key (e.g. *z,z,z,m,m,m,z,z,z,m,m,m*, or *z,m,z,m*), they were excluded. Finally, participants were excluded and replaced during the testing phase if they did not comply with all instructions throughout the experiment or if there was any misfunctioning in the stimulation equipment during the session. This also included if participants reported any discomfort from the stimulation or if the Nurostym device identified that the electrode impedances were too high and self-terminated the stimulation. Participants who meet any of the above exclusion criteria were removed before the experimenter was unblinded to the data and they were replaced during the data collection phase, as outlined above.

### Behavioural assessments

#### Finger-Tapping Random-Sequence Generation Task (FT-RSGT)

The FT-RSGT required participants to respond in a random sequence to an ongoing metronome tone by pressing one of two response-buttons at a time (see Figure 3). This task was selected to replicate the methodology used by Rasmussen et al. (2024) and because it was designed to be a more reliable test for detecting periods of mind wandering due to the large number of trials and more sensitive measures of task performance (Alexandersen et al., 2022; Boayue et al., 2021). The two response buttons corresponded to two separate keys (‘z’ for left-hand and ‘m’ for right-hand). The metronome tone was presented at 440Hz for 75ms, with a 750ms inter-stimulus interval and participants were instructed to time their taps with the tone as accurately as possible. During the task, participants were asked to maintain their focus on a white (RGB 255 255 255) fixation cross in the centre of the screen with a grey (RGB 128 128 128) background.

**Figure 3.**
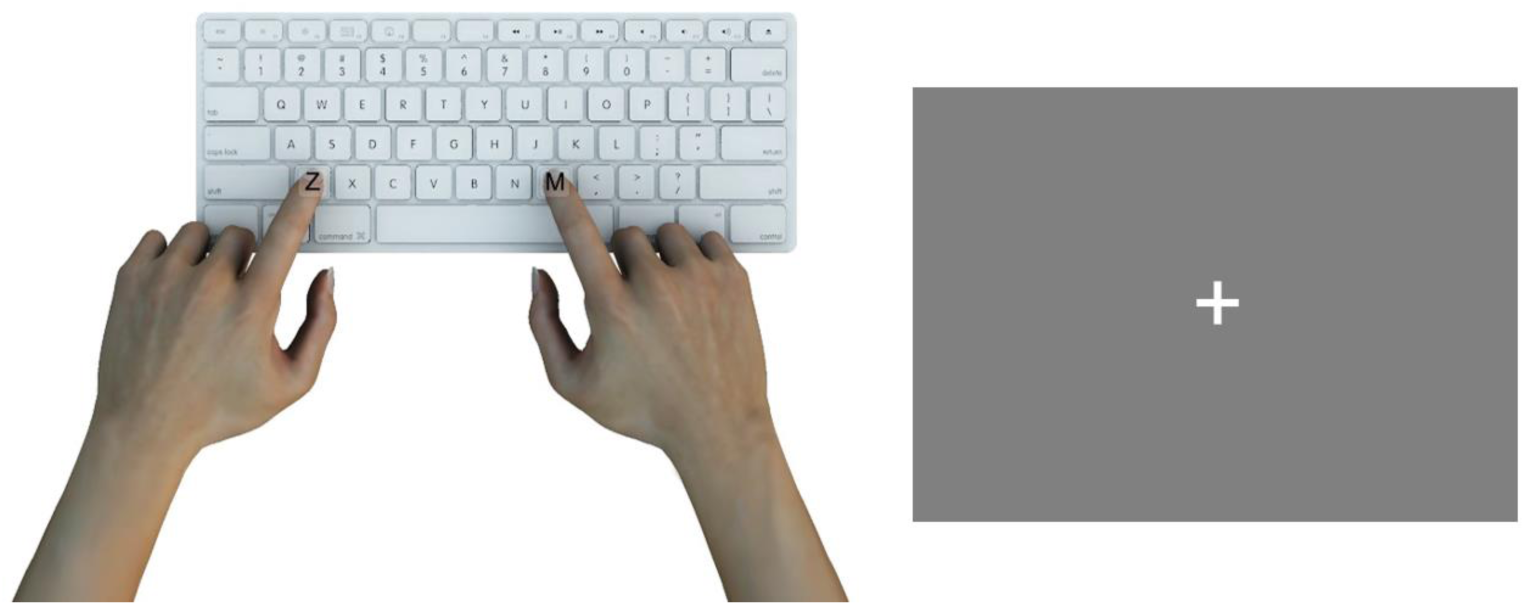
Finger-Tapping Random-Sequence Generation Task. Illustration of the two keys participants used to generate the random sequences (on the left) and the display screen (on the right).

The task was presented on a 24-inch LED monitor, with a refresh rate of 100 Hz. Participants sat approximately 70cm away from the monitor and used a standard Macintosh keyboard and mouse to respond. The auditory tone was also presented through CREATIVE GigaWorks T40 Series II speakers. There were initially 20 practice trials in the training block, which could be completed twice to consolidate participants understanding of the task. Participants then completed training on the thought probes, followed by another 20 practice trials, including an example of the four thought probes after the final trial. There was then a baseline block that consisted of 720 trials, running for approximately 10 minutes, before participants were administered with levodopa or a placebo and then waited 40-minutes preceding the stimulation block. The stimulation block included 2160 trials, and ran for approximately 30 minutes, including a 30 second break after approximately 15 minutes.

There were two measures of task performance: the randomness of the sequences and participants variability in the timing of their responses. The randomness of participants sequences was determined by a measure of approximate entropy, which is designed to calculate the predictability of the next item in a sequence, based on a specified number of previous items, *m*. This study employed an *m* = 2, which replicated the *m* value used by Rasmussen et al. (2024). This measure was selected as there is evidence for a relationship between executive functioning and randomness, such that directing more executive resources towards the FT-RSGT, will result in more random sequences (Boayue et al., 2021). Thus, a smaller ApEn score indicated more repetitive patterns in the data and a larger score represented greater randomness in the patterns, which can be used to infer participants were more focused on the task. Participants performance was also measured through behavioural variability, which is the deviation in their responses from the metronome tone. This was calculated using the 20 trials prior to each set of mind wandering probes, where the standard deviation of the difference between the tone and response for these trials was included in the analyses.

### Mind wandering probes

There were also four thought probes presented throughout the task, which were designed to assess the contents of participants thoughts. The four probes appeared together each time and they were pseudo-randomly presented every 45 to 75 seconds during the baseline and stimulation blocks. Thus, there was a total of 10 probes in the baseline block and 30 probes in the stimulation block. The four questions were: (1) Before the probe, were you thinking about something other than the random sequence generation task; (2) Before the probe, was your mind wandering around freely; (3) Were you actively directing your thoughts; and (4) Was your mind stuck on something. The questions were designed to ask participants about their thoughts in the 10-15 seconds before the probes were presented. Participants responded to each question on a 7-point Likert scale which ranged from “Not at all” (1) to “Very much” (7), with the middle point (4) labelled “Moderately”. These responses were made using to 1-7 keys on the keyboard, and there were no time constraints to respond.

Participants were trained on the four thought probes at the beginning of the session, going through detailed explanations of the four types of thoughts, alongside example scenarios where they could occur. These explanations were based on Kam et al. (2021), however they were designed to replicate Rasmussen et al. (2024). Participants were then tested on their understanding of the four questions by explaining their responses for each thought probe in the context of four example scenarios. The full description of the probes and examples can be found in the supplementary materials. The thought probe information was presented in the centre of a grey background (RGB 128 128 128) in white Arial font (visual angle = 1.1°).

### Levodopa protocol

Each participant began the session with their blood pressure and heart rate being measured. After training on the task and a baseline block of the FT-RSGT, they then received either levodopa (Madopar® 125 tablet: Levodopa 100mg/ Benserazide Hydrochloride 25mg) or a placebo tablet (Centrum® for women multivitamin)^1^. The Madopar table combines levodopa and benserazide hydrochloride to prevent the immediate uptake of the drug, as the benserazide component is unable to cross the blood-brain barrier which inhibits the early conversion of levodopa to dopamine (Contin & Martinelli, 2010). To double blind the main experimenter and participant to the drug condition, an additional experimenter who, was not otherwise involved in the testing sessions, oversaw the drug protocol. Participants were then required to wait approximately 40 minutes after the drug administration before completing the stimulation block, to ensure the task was undertaken when the plasma concentration was around its peak (Contin & Martinelli, 2010). At the end of the session, participants and the experimenter were asked to select which dopamine drug manipulation group they were in (levodopa, placebo), to assess the efficacy of the drug manipulation blinding. They were again assessed on their confidence in this decision, by asking “How confident are you in your judgement of your dopamine drug condition?”. The responses were rated on a 7-point Likert scale, ranging from “Not at all confident” (1) to “Extremely confident” (7), with the midpoint as “Moderately confident” (4). This data was used to compare the proportion of correct guesses across both conditions, whereby a lower proportion of correct guesses across the two groups indicated that the blinding was effective for these conditions.

### Stimulation protocol

Each participant received 2mA anodal or sham HD-tDCS over the left PFC. The stimulation was delivered online, in conjunction with the FT-RSGT, with each participant receiving one of two stimulation protocols in conjunction with either the levodopa or placebo drug (4 groups in total). Whether stimulation was active or sham was double-blinded. At the end of the session, participants and the experimenter were asked whether they received active or sham stimulation, to assess the effectiveness of the blinding. Participants and the experimenter were also asked to rate their confidence in their decisions, asking, “How confident are you in your judgement of your stimulation condition?”. This question consisted of a 7-point Likert scale, ranging from “Not at all confident” (1) to “Extremely confident” (7), with the mid-point labelled as “Moderately confident” (4).

### HD-tDCS montage

The electrode placement was determined using the International 10-20 EEG system, with the anode placed over F3 and the four surrounding reference cathodes at F7, Fp1, C3 and Fz (see Figure 4). This was designed to replicate the PFC montage used by Rasmussen et al. (2024). The HD-tDCS was administered at 2.0mA current intensity, with 0.5mA to each of the four reference electrodes, respectively. The stimulation was delivered using a Nurostym stimulator, with a 4 x 1 ring electrode arrangement. The electrodes were 12mm Ag/AgCl electrodes which were secured to the scalp using a cap and conductive gel. The stimulation lasted for 30 minutes, including a 30 second ramp up and down period. The sham stimulation was delivered for 75 seconds, with the same ramp up and ramp down times. All participants first completed a 10-minute baseline block of the FT-RSGT, before completing 30-minutes of 2mA active or sham stimulation in conjunction with the task.

**Figure 4.**
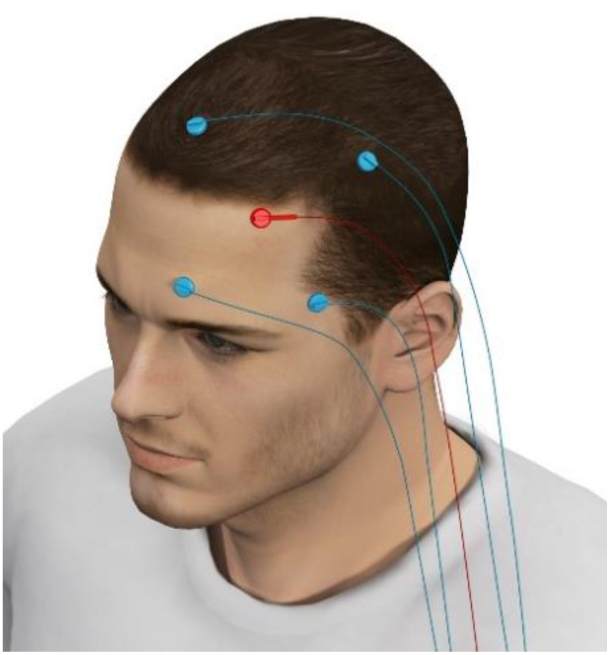
HD-tDCS montage. The PFC arrangement, with the target electrode at F3 and the surrounding four cathodes.

### Self-report questionnaires

Given the FT-RSGT requires participants to respond accurately in time to a metronome tone, it was important to account for any influence of video game or musical training on participant’s response variability. Thus, at the beginning of the session, participants were asked how many hours they spent playing video games and musical instruments each week, and this information was entered into the randomisation script to ensure these two variables were balanced across the four groups (see supplementary materials). Measures of heart rate, blood pressure and mood were also taken at the beginning and at the end of the session. Mood was assessed via the Bond-Lader Visual Analogue Scale (BL-VAS; Bond & Lader, 1974), which is a sixteen-item scale, with each element rated on a 10-point analogue scale. The total score was calculated across all the items.

During the HD-tDCS electrode set up, participants were also given two tests, designed to measure trait impulsivity and participants’ morningness and eveningness predisposition. This is because there is evidence to suggest that individual’s attentional impulsiveness is correlated with variation in dopamine D2 and D3 receptor availability (Buckholtz et al., 2010; Taylor et al., 2018). Furthermore, individuals’ circadian arousal rhythms, measured via their disposition as a morning or evening person, have also been found to be correlated with cognitive control functions (Anderson et al., 2014) and an individual’s propensity to mind wander (Carciofo et al., 2013, 2014). The Barratt Impulsivity Scale (BIS-11) is a 30-item questionnaire, which rates each item on a 4-point Likert scale from “Rarely/Never” (1) to “Almost always/always” (4). It is designed to assess three factors which predict impulsive personality traits: attentional (attention and cognitive instability impulsiveness), motor (motor and perseverance), and non-planning (self-control and cognitive complexity). The Morningness-Eveningness questionnaire (Horne & Ostberg, 1976) was used to assess participants activity and alertness across different times of day. There are 19 items, consisting of both Likert and time-scale questions, which were totalled to obtain a global score. This was converted into a 5-point scale, designed to categorise the different circadian rhythms across participants.

In addition, mind wandering has been found to be associated with individual’s trait mindfulness abilities and with pervasive negative thoughts, or ruminative thoughts (Jonkman et al., 2017; Mrazek et al., 2012). Furthermore, there is evidence to suggest that ADHD symptomology is predictive of mind wandering (Franklin et al., 2017; Jonkman et al., 2017) and deficits in dopamine concentrations have also been found in clinically diagnosed ADHD patients (Mehta et al., 2019). Thus, there were three questionnaires given at the end of the session to control for any individual differences in these variables between the experimental conditions. These questionnaires included the Mindful Attention and Awareness Scale (MAAS), the Rumination Response Scale and the Adult ADHD Self-Report Scale (see supplementary materials for questionnaire details). The MAAS assessed how present participants were in their daily lives via 15 items which were rated on a 6-point scale ranging from “Almost Always” (1) to “Almost Never” (6). Participants scores were calculated via the mean of the 15 items, with higher levels of mindfulness being represented by higher scores. The Rumination Response Scale was included to assess how often participants engaged in different ruminative thoughts when they were feeling down, sad, or depressed. Participants rated each of the 22 items on a 4-point Likert scale from “Almost never” (1) to “Almost always” (4) and the scores were added together. Greater ruminative thoughts were represented by higher total scores. Finally, the Adult ADHD Self-Report Scale is made up of 18 questions which are designed to capture the frequency that participants exhibit symptoms associated with ADHD. All questions were rated on a 5-point Likert scale, ranging from “Never” (1) to “Very Often” (5), with the total score calculated by totalling all the ratings together and higher scores indicated a greater number of ADHD tendencies.

The end of session questionnaire also included a detailed assessment of participants involvement with video games and musical instruments, alongside questions addressing participants perspective on their task performance and their experience with the drug and stimulation. Finally, participants were asked to rate their motivation to complete the task on a 7-point Likert scale ranging from “Not at all motivated” (1) to “Extremely motivated” (7). This question asked participants, “How motivated were you to perform well in this task?” (Seli et al., 2015). Motivation levels have been linked to task performance outcomes (Brosowsky et al., 2020) and there is evidence to suggest dopamine levels are linked to motivation (Mohebi et al., 2019), thus it was important to assess the motivation levels between the four groups to ensure there were no differences which may be influencing the results. For the full end of session questionnaire, refer to the supplementary materials.

### Analysis Overview

#### Overview

At the end of each participants session, all raw data files were uploaded to The University of Queensland Research Data Manager cloud storage. All analyses were completed using a combination of JASP and RStudio, specifically employing the *brms* (Bayesian Regression Models using Stan; Bürkner, 2017) and *BayesFactor* packages (Mulder et al., 2021). There is currently no previous research investigating the effect of dopamine concentrations and stimulation on individuals’ propensity to mind wander, thus this study employed a default Jeffreys-Zellner-Siow prior of r = 0.707, centred around 0 (Rouder et al., 2009). All analyses assessed whether the results were more likely under the null or alternative model, which were interpreted by BF_10_ and BF_01_. Here, we used a conservative approach to interpret BF values, with BF_10_ > 6 and BF_01_ > 6 taken to represent meaningful evidence for an alternative or null effect, respectively (Lee & Wagenmakers, 2013). The analyses primarily focused on the effects of stimulation and dopamine on freely moving thought and task unrelated thought, as these thought types had previously been shown to be modulated by tDCS applied to the left PFC (Boayue et al., 2021; Filmer et al., 2019; Rasmussen et al., 2024), however the effects on deliberately and automatically constrained thought were also investigated in a more exploratory manner. All the raw data, study materials, supplementary materials and results, and analysis scripts from our study can be located at https://doi.org/10.48610/e9cbbaf.

#### Post-study data exclusion

The post-study exclusion criteria for participants replicated the criteria used by Rasmussen et al. (2024) and was employed once data collection was completed, thus these participants were not replaced in the final sample. Firstly, across all participants in the baseline block, individuals who scored more than 3 standard deviations above or below the sample mean for their approximate entropy and behavioural variability scores were excluded from the study. This resulted in five participants being removed. Furthermore, in the stimulation block, participants were removed if they scored more than 3 standard deviations above or below their respective group’s mean for their approximate entropy or behavioural variability scores, or their mean responses to any of the four thought probes. This resulted in an additional four participants being removed, creating a final sample of 231 participants. Finally, to ensure that extreme outliers during the task did not skew any time on task effects, individual trials which were greater than 3 standard deviations above or below the mean for each group’s approximate entropy and behavioural variability scores were also be removed from the analyses. This resulted in 63 trials (1.37% of trials) being removed from the active and sham groups, across the placebo condition; 65 trials (1.45% of trials) being removed from the levodopa and placebo groups, across the sham condition; and 76 trials (1.65% of trials) being removed from the active levodopa and active placebo groups. This resulted in a final trial sample of 4563, 4494 and 4604 for each analysis, respectively.

#### Applying modelling to investigate the dynamic thought types

This study primarily employed hierarchical order probit modelling to investigate the effects of HD-tDCS and dopamine on the dynamic thought types (Alexandersen et al., 2022; Boayue et al., 2021; Rasmussen et al., 2024). This analysis technique treated the thought probe responses as ordinal data, which allowed for investigation into time on task effects and individual’s response variability across the duration of the task (Rasmussen et al., 2024). Participants thought probe responses were entered as the dependent variable into the models and there were several predictor variables, relating to the measures of task performance (behavioural variability and approximate entropy), block and trial data, alongside their interactions, however participants stimulation condition (2mA active vs sham) or dopamine condition (levodopa vs placebo) was entered in as the key predictor for the respective analyses. The specific predictors employed in each probit model are explained in detail below. The model weights were interpreted using two methods. The first method was by calculating the pareto smoothed importance sampling leave-one-out cross-validation scores (PSIS-LOO; Vehtari et al., 2017, 2022) and then comparing the LOO information criterion values (LOOIC; Vehtari et al., 2017) using a stacking procedure (Vehtari & Gabry, 2018; Yao et al., 2018). The second method was applying pseudo-Bayesian model-averaging (pseudo-BMA; Vehtari & Gabry, 2018; Yao et al., 2018).

In analyses where the LOOIC and pseudo-BMA winning models do not agree on a preferred model, the winning LOOIC model was interpreted, as this method has been found to have greater predictive accuracy when the true model is not in the model list, by selecting the model with the best predictive distribution (Vehtari & Gabry, 2018; Yao et al., 2018). This is consistent with the analysis practices used by Rasmussen et al. (2024). Where there were inconsistencies in model interpretations, they may be driven by differences in the amount of data for the baseline (10 probe sets) and stimulation (30 probe sets) phases. Thus, where needed to address model discrepancies, the model comparisons were re-run on the stimulation data alone.

#### The effect of HD-tDCS on task unrelated and freely moving thought

We hypothesised that 2mA anodal HD-tDCS to the left PFC would reduce freely moving thought, relative to the sham condition (H_1a_). We also hypothesised that 2mA anodal HD-tDCS would reduce task unrelated thought, relative to the sham condition (H_1b_). We first employed the hierarchical order probit modelling to investigate the effect of anodal stimulation on both thought types, alongside the effects of stimulation on task performance. This consisted of comparing 23 models, increasing in complexity, which included the following predictor variables and their interactions: stimulation (active vs sham), behavioural variability, approximate entropy, trial, and block (baseline vs stimulation). There was one probit model assessing the effects on freely moving thought, and the other assessing the effects on task unrelated thought.

In addition, to investigate the direct effect of stimulation on both freely moving and task unrelated thought, we employed a Bayesian independent samples t-test for both thought types. The t-test compared the active anodal HD-tDCS group to the sham group, for participants who were in the placebo drug condition. While tests were conducted for both thought probes, the results from the freely moving thought t-test was one of the two tests used for the stopping rule in this study, as there was evidence for 2mA anodal stimulation reducing freely moving thought found by Rasmussen et al. (2024).

#### The effect of dopamine on task unrelated and freely moving thought

As there is evidence hinting at a relationship between dopamine and mind wandering (Cools, 2008; O’Callaghan et al., 2021), we were also interested in investigating the direct effects of dopamine on freely moving and task unrelated thoughts. We hypothesised that there would be an effect of levodopa, relative to the placebo drug, on freely moving and task unrelated thought (H_2_). To assess these effects, we used two hierarchical order probit models – one with freely moving thought as the outcome variable and the other assessing the effects on task unrelated thought responses. Each probit model consisted of 23 models of increasing complexity, which included the following predictor variables and their interactions: pharmacological manipulation (levodopa vs placebo), behavioural variability, approximate entropy, trial, and block (baseline vs stimulation). For both thought types, we also assessed the direct effect of dopamine by applying a Bayesian independent samples t-test, comparing the levodopa condition to the placebo condition, for participants who were allocated to the sham group.

#### The interactive effects of dopamine and stimulation on the dynamic thought types

Given findings that suggest the effects of tDCS on behavioural outcomes may be mediated by individuals’ dopamine concentration (Leow, Marcos, et al., 2023), this study predicted there would be a difference in the effect of 2mA HD-tDCS, in combination with levodopa on freely moving thought and task unrelated thought, relative to the active stimulation with placebo group (H_3_). This was first analysed using hierarchical order probit modelling to assess the combined effect of stimulation and dopamine. There was a probit model run for both freely moving and task unrelated thought, with 23 models included in each. Specifically, they included the key predictor as: dopamine-stimulation manipulation (active stimulation with levodopa vs active stimulation with placebo), alongside the additional predictors – behavioural variability, approximate entropy, trial, and block (baseline vs stimulation).

Furthermore, there was a Bayesian independent samples t-tests employed for both freely moving and task unrelated thought, to assess the effect of stimulation in conjunction with dopamine. They compared the active HD-tDCS with levodopa condition to the active HD-tDCS with placebo condition for each thought type. Importantly, the freely moving thought t-test was also used to determine the stopping rule for this study. This test was selected, as we believed a meaningful interaction between stimulation dosage and dopamine would contribute to our understanding the neurochemical mechanisms underlying mind wandering.

#### Control analyses

The application of levodopa can occasionally result in side effects including nausea (Chen et al., 2020) and changes in mood state (Beaulieu-Boire & Lang, 2015). Thus, to ensure that there were no changes in blood pressure, mood, or heart rate due to the drug administration, we ran three 2 (Time: before drug administration, ∼2 hours after drug administration) x 2 (Drug: levodopa, placebo) Bayesian between-subject ANOVAs, with blood pressure (diastole and systole)^2^, BL-VAS scores and heart rate as the dependent measures, respectively. We interpreted a BF_excl_ > 6 for the interaction term as moderate evidence for there being no meaningful differences between the groups and thus the null hypothesis was accepted.

#### Testing for baseline differences

To investigate any group differences in participants’ responses in the self-report measures, we used five one-way between-subjects ANOVAs. This included responses for the BIS-11, the Morningness-Eveningness questionnaire, the Adult ADHD Self-Report Scale, the Rumination Response Scale, and the MAAS, as the respective dependent measure for the five ANOVAs. The four stimulation conditions were also included as the predictor variable in each analysis (sham HD-tDCS with placebo; sham HD-tDCS with levodopa; anodal HD-tDCS with placebo and anodal HD-tDCS with levodopa). If there were any meaningful differences found between the groups for any ANOVA, these would be followed up using t-tests, and any measures with meaningful group differences would also be included as a covariate in the modelling analyses.

To ensure there were no differences between the groups in their baseline responses to the thought probes, we also reran the 2 (Stimulation: active, sham) x 2 (Drug: levodopa, placebo) Bayesian between-subjects ANOVA on the baseline data alone, for both freely moving thought and task unrelated thought. We interpreted a BF_excl_ > 6 for the interaction term as moderate evidence for there being no meaningful differences between the groups and thus the null hypothesis was accepted.

#### Assessing blinding

We assessed the effectiveness of the blinding for the stimulation and pharmacological conditions that participants were allocated to. For each of these factors, we assessed the proportion of correct guesses between the two groups – the active and sham stimulation conditions; and the levodopa compared to placebo conditions. This was assessed for both the participants and the experimenters.

#### Exploratory investigation into deliberately and automatically constrained thought

In addition to assessing the effects of HD-tDCS and dopamine on freely moving and task unrelated thought, we also assessed these effects on deliberately constrained and automatically constrained thought. As there was no evidence to date for these thought types being modulated via tDCS to the PFC (Rasmussen et al., 2024), these analyses were exploratory. We ran the same hierarchical order probit analyses used in the central analyses above, employing 23 models of increasing complexity, and the key predictor alternated between stimulation, and drug, to assess these effects on each thought type. We also included the following predictors and their interactions in all the probit models: behavioural variability, approximate entropy, trial, and block (baseline vs stimulation). However, to investigate the effect of stimulation, we employed one probit model for each thought type, including stimulation (active vs sham) as the additional predictor. In addition, we ran two models, one for both thought types, which included the dopamine condition (levodopa vs placebo) as the added key predictor. Finally, to investigate the interactive effect of stimulation and dopamine we ran one model for both deliberately and automatically constrained thoughts. This pair of models included active stimulation with levodopa vs active stimulation with placebo as the key predictor.

In addition to the modelling analyses, we employed the same Bayesian independent samples t-tests which were applied in the main analyses. Each t-test was run for both deliberately constrained thoughts and automatically constrained thoughts. We first compared the anodal HD-tDCS group to the sham group, for participants who were in the placebo drug condition. We then assessed the effect of dopamine alone, by comparing the levodopa group to the placebo group, for participants in the sham condition. Finally, we ran a t-test for each thought type, to assess the combined effect of stimulation and dopamine on the frequency of each thought probe. Each compared active HD-tDCS with levodopa to active HD-tDCS with placebo.

## Results

### The effect of HD-tDCS on task unrelated and freely moving thought

To investigate the effect of 2mA anodal HD-tDCS, applied to the left PFC, on freely moving thought, we employed hierarchical order probit modelling. The probit modelling included 23 models of increasing complexity which were weighted using two model selection procedures, and the winning LOOIC model for each probit model was chosen for this study (refer to Figure S1 for the model comparisons for each set of probit models). This analysis technique was used to replicate the approach used by Rasmussen et al. (2024), who found preliminary evidence that anodal stimulation, relative to sham stimulation, reduced freely moving thought in this region. Thus, we hypothesised anodal HD-tDCS would reduce freely moving thought, relative to the sham group, across participants in the placebo drug condition (H_1a_). However, we found no evidence to support this hypothesis. The winning model did not include the key predictor to demonstrate an effect of stimulation on freely moving thought, which was a stimulation x block interaction. Furthermore, the Bayesian independent samples t-test found anecdotal evidence against an effect of stimulation on this thought type (BF_01_ = 2.60), although this result did not reach our threshold to be considered meaningful (BF_10_ > 6 or BF_01_ > 6). The full output from these analyses can be found in the supplementary results: the top LOOIC and Pseudo-BMA model selection weights for the complete datasets and stimulation block datasets are in Table S1 and S2, respectively. The predictors for each preferred LOOIC model are in Figure S2 and Table S3.

We also hypothesised 2mA anodal HD-tDCS would reduce task unrelated thought, relative to sham stimulation, across the placebo groups (H_1b_). However, we again, found no evidence to support this hypothesis. The probit modelling demonstrated no evidence for a meaningful block x stimulation interaction, nor a main effect of stimulation (*b* = .08, 95% CI [−.20, .35] and *b* = .16, 95% CI [−.23, .53], respectively; see Figure 5). Interestingly, the model did include a meaningful positive behavioural variability x stimulation interaction (*b* = .65, 95% CI [.21, 1.09]). This effect was conditional on participants self-reported state, such that when participants in the active stimulation group reported task unrelated thoughts, they also had greater variability in their responses compare to participants in the sham stimulation group. This suggests active stimulation may impair behavioural performance to a greater extent than sham stimulation when participants were experiencing task unrelated thoughts. However, as the stimulation variable includes both the baseline and stimulation block data, this is not a true measure of the effect of HD-tDCS on task performance. In the stimulation block data alone, no stimulation effects were selected in the winning model and the t-test also found strong evidence against an effect of HD-tDCS on task unrelated type (BF_01_ = 10.12). While these findings do not support our hypothesis, they do align with a recent high powered registered report, using the same 2mA HD-tDCS montage, finding no evidence for this effect (Alexandersen et al., 2022).

**Figure 5.**
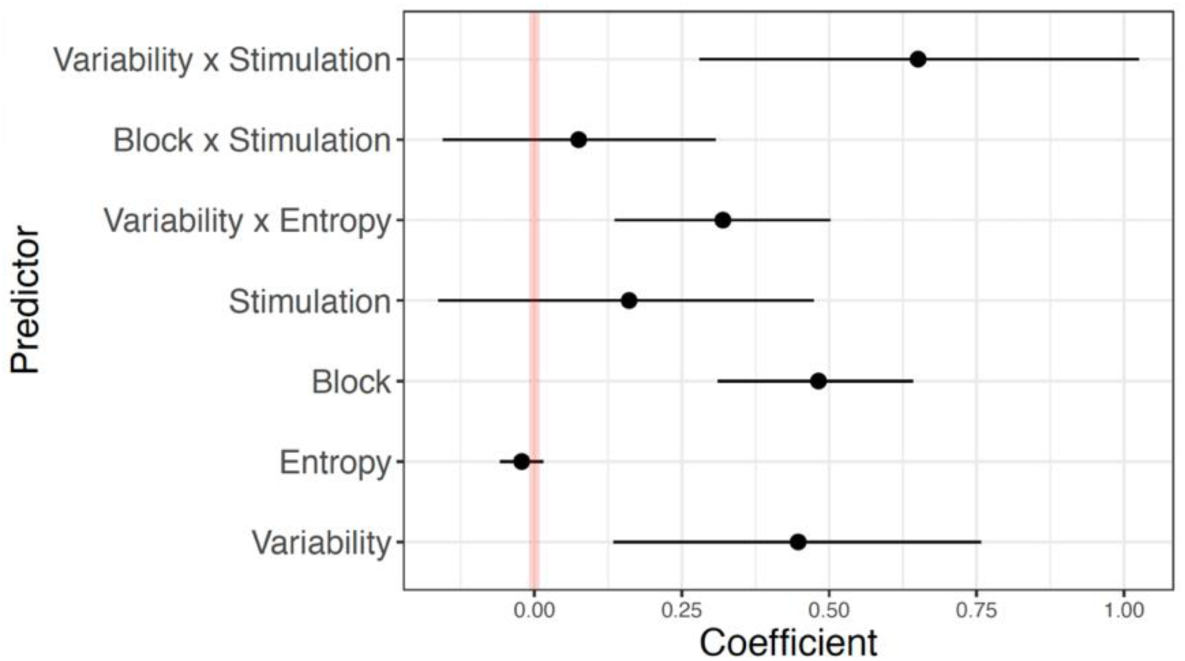
Winning probit model for the effect of stimulation on task unrelated thought. The winning complete model for the effect of HD-tDCS, across the placebo drug condition, on task unrelated thought. This shows no effect of stimulation on task unrelated thought, however there is a meaningful interaction between behavioural variability and stimulation.

### The effect of dopamine on task unrelated and freely moving thought

We were also interested in understanding the role of dopamine in modulating freely moving and task unrelated thought, given evidence suggesting there may be a relationship between dopamine and mind wandering (Cools, 2008; O’Callaghan et al., 2021). We predicted there would be a difference in the effect of levodopa, relative to the placebo group, on freely moving thought, across the sham condition (H_2a_). There was evidence that levodopa may reduce freely moving thought. The winning model found a negative main effect of levodopa (*b* = −.42, 95% CI [−.72, −.11]), which suggests levodopa reduced freely moving thought, relative to the placebo condition (see Figure 6). However, this variable included the baseline and stimulation block data, which means it cannot be used to determine the effect of levodopa alone. The effect of levodopa on freely moving thought was reinforced by evidence from the stimulation block alone modelling, which found levodopa reduced freely moving thought relative to the placebo condition, via a meaningful negative main effect of levodopa (*b* = −.41, 95% CI [−.74, −.10]). Given this effect is specific to responses after the levodopa was administered, it strengthens the reliability of this finding. Furthermore, as discussed below, there is moderate evidence from the Bayesian between-subjects ANOVA for there being no baseline differences between the experimental groups (BF_excl_ = 4.91) for freely moving thought and while this does not reach our threshold to be considered meaningful, it suggests baseline differences are not driving the main effect of levodopa found in the complete dataset. The validity of the finding is also reinforced as there are only 10 probes in the baseline block and 30 probes in the stimulation block, thus, it is unlikely the baseline block could drive the main effect found across blocks.

**Figure 6.**
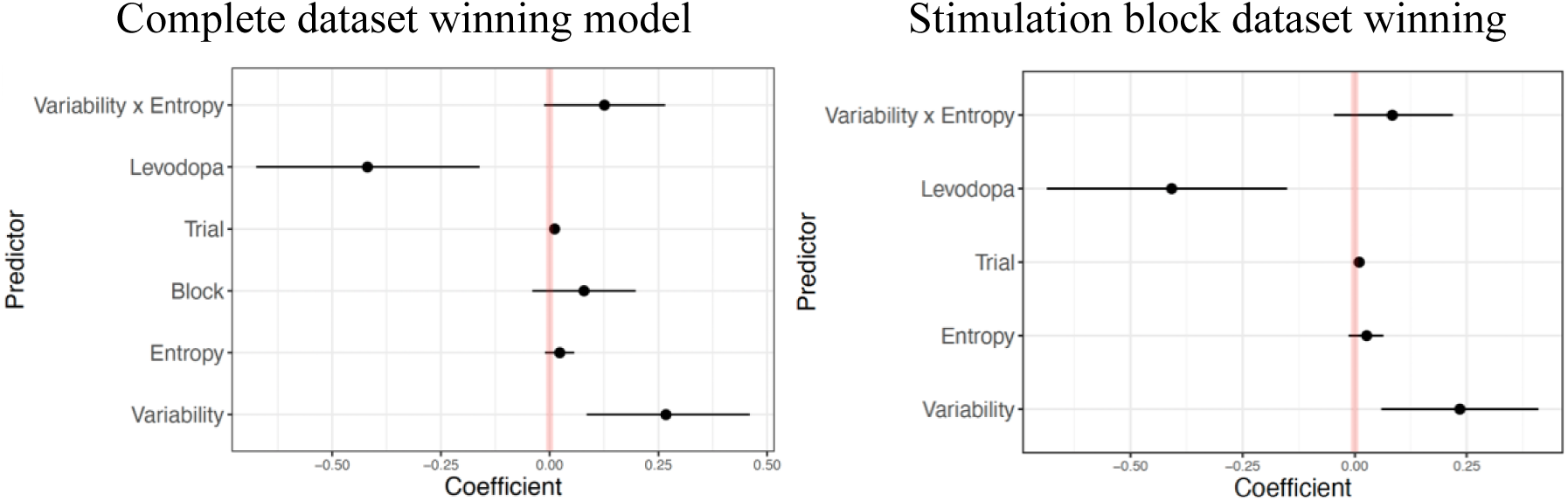
Winning probit models for the effect of dopamine on freely moving thought. The effect of levodopa vs placebo, across the sham stimulation condition, on freely moving thought. The left image displays the winning LOOIC model for the complete dataset (including the baseline block) and the right image displays the winning LOOIC model for the stimulation block alone, with both models finding levodopa reduced freely moving thought.

Finally, the t-test found anecdotal evidence for a difference between the levodopa and placebo conditions (BF_10_ = 2.64), however this did not reach the threshold to be considered meaningful. Overall, these findings provide evidence that levodopa may reduce freely moving thoughts relative to the placebo drug.

We also predicted there would be a difference in the effect of levodopa, relative to the placebo group, on task unrelated thought, across the sham conditions (H_2b_). However, there was no evidence for any effect of dopamine on this thought type. Both the key block x levodopa interaction and a main effect of levodopa were not meaningful in the winning model (*b* = .09, 95% CI [-.21, .39] and *b* = −.237, 95% CI [−.68, .13], respectively). This was consistent with the stimulation block alone, which found no evidence for a main effect of levodopa (*b* = −.20, 95% CI [−.57, .20]). Finally, the Bayesian independent samples t-test found moderate, but not meaningful, evidence for a null effect of dopamine on task unrelated thought (BF_01_ = 3.28).

### The interactive effects of dopamine and stimulation on the dynamic thought types

Given there is evidence to suggest that the effects of levodopa may interact with tDCS to affect behavioural outcomes (Curtin et al., 2024; Fresnoza et al., 2014; Leow, Marcos, et al., 2023), we hypothesised that there would be a difference between the effect of active stimulation with and without levodopa, on freely moving thought (H_3a_). However, there was no evidence for this effect. There were no levodopa-stimulation predictors selected in the winning probit model and the t-test also found moderate evidence for no differences between these groups (BF_01_ = 4.84), however this did not meet our threshold to be considered meaningful. Overall, these findings do not support our hypothesis, as they suggest the combination of levodopa and HD-tDCS did not affect freely moving thought.

We also hypothesised that there would be a difference between the effect of active stimulation, with and without levodopa, on task unrelated thought; however, there was no evidence for this effect (H_3b_). Specifically, there was no evidence for the block x levodopa-stimulation interaction, nor for the main effect for the levodopa-stimulation predictor (*b* = .06, 95% CI [-.26, .38] and *b* = −.05, 95% CI [−.42, .31], respectively; see Figure 7). While there was no direct effect of levodopa and stimulation on this thought type, the model did find a meaningful negative behavioural variability x levodopa-stimulation interaction (*b* = −.54, 95% CI [−.96, −.14]). This suggests that when the levodopa and active stimulation were combined participants had less variability in their scores during periods of task unrelated thought, than when the active stimulation was delivered alone. However, this interaction includes the baseline data and thus it is not a true measure of the effect of HD-tDCS combined with levodopa on behavioural variability. The winning model for the stimulation block alone was consistent with the complete dataset, finding no main effect for the levodopa-stimulation predictor on task unrelated thought (*b* = .06, 95% CI [-.30, .42]). Finally, the t-test found moderate, but not meaningful, evidence for a null effect (BF_01_ = 4.94). Together, these findings suggest that our hypothesised interaction between dopamine and stimulation affecting task unrelated thought was not supported.

**Figure 7.**
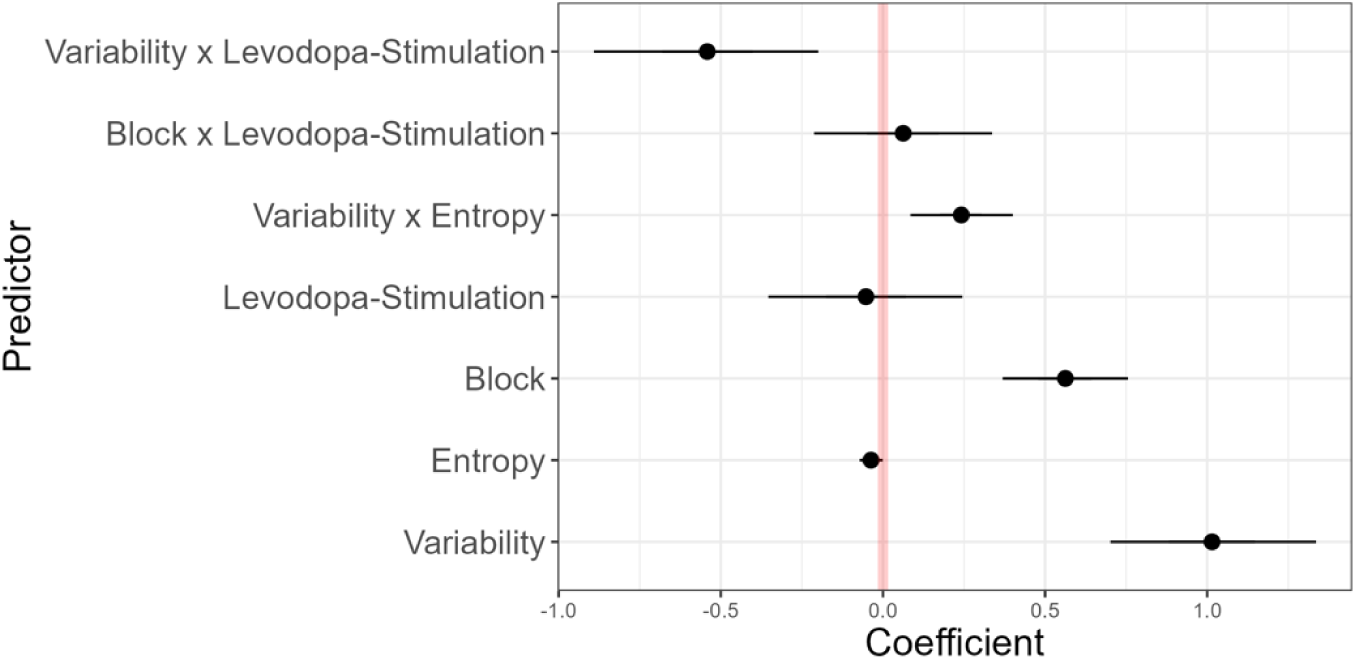
Winning probit model for the combined effect of dopamine and stimulation on task unrelated thought. The winning complete model for the combined effect of HD-tDCS and levodopa on task unrelated thought. This shows no interactive effect of stimulation and dopamine, however there is a meaningful interaction between the stimulation-dopamine predictor and behavioural variability.

### Control analyses

To assess any psychological changes due to the administration of levodopa, including changes in blood pressure and mood, we ran Bayesian between-subject ANOVAs with systolic and diastolic blood pressure, BL-VAS scores and heart rate as the dependent measures and Time (pre-drug administration, ∼2 hours after drug administration) x Drug Condition (levodopa, placebo) as the independent measures. We predicted there would be no change in heart rate, blood pressure or mood with the administration of levodopa. Overall, the findings supported this hypothesis. The diastolic blood pressure Time x Drug Condition interaction was the only measure to reach our threshold to be considered meaningful, providing moderate evidence against an effect (BF_excl_ = 6.56). Heart rate (BF_excl_ = 4.21) and mood (BF_excl_ = 3.86) also provided moderate evidence for excluding the interaction term, however this did not reach our threshold to be considered a meaningful null effect. There was anecdotal evidence for an effect on systolic blood pressure, as shown by the Time x Drug Condition interaction (BF_incl_ = 2.10). While this did not reach the threshold for a meaningful effect, it suggests that systolic blood pressure may have been reduced to a greater extent for participants in the levodopa condition (mean reduction in systolic blood pressure, 4.66), compared to the placebo condition (mean reduction in systolic blood pressure, 1.57). Please refer to Table S4 for a complete summary of the pre- to post-changes for each physiological measure.

### Testing for baseline differences

To investigate any group differences in participants self-report measure responses, we ran five one-way between-subjects ANOVAs which included the responses from the BIS-11, the Morningness-Eveningness questionnaire, the Adult ADHD Self-Report Scale, the Rumination Response Scale and the MAAS as the respective dependent measure and the four groups as the independent variable (active or sham stimulation combined with either the levodopa or placebo drug). We hypothesised there would be no differences in the relevant traits between the four groups, which was supported by the results. There was moderate to strong evidence for excluding the group variable in all five questionnaires (BF_excl_ > 8.78 for all), which indicates there were no differences for the self-report measures between the experimental groups.

We also assessed whether there were any group differences in participants baseline responses to the freely moving thought and task unrelated thought probes. There was moderate evidence against any group differences for task unrelated thought, shown by BF_excl_ = 9.39 for the group variable, and thus the null hypothesis was accepted. There was also moderate evidence against any group differences for freely moving thought (BF_excl_ = 4.91), however this value did not reach the BF_excl_ > 6 threshold to be considered meaningful.

### Assessing blinding

It was also important to assess the effectiveness of the blinding for both the stimulation and pharmacological conditions that participants were allocated to, by assessing the proportion of correct guesses between the two groups (i.e. active vs sham and levodopa vs placebo). To assess whether these effects were meaningful, we also ran a Bayesian independent samples t-test, which can be found in the exploratory section below as this additional analysis was not included at Stage 1. We found evidence to support our hypothesis that both participants and the experimenter would be unable to correctly identify their stimulation group. Overall, the stimulation blinding appeared to be effective for the participants, as 113 out of 231 participants correctly guessed their group (49%). There was a bias towards participants selecting the active stimulation condition, regardless of their group allocation, as 76 out of 117 participants in the active group correctly guessed their condition (65%) and 41 guessed incorrectly (35%). In the sham group, only 37 participants correctly guessed their group (32%) and 77 out of 114 participants guessed incorrectly (68%). The experimenter blinding had the opposite trend, whereby they were biased towards the sham condition, regardless of group and this may be due to a more conservative approach towards claiming that participants received active stimulation (e.g. by accounting for the participants verbal or visible response to the start of stimulation). Overall, the experimenter correctly classified the stimulation group of 133 out of 231 participants (58%). However, they only classified 53 out of 117 participants correctly in the active condition (45%) and incorrectly classified 64 participants (55%). In the sham condition, the experimenter correctly classified 80 out of 114 participants (70%) and incorrectly classified 34 participants (30%). These findings suggest that both the participant and experimenter blinding for stimulation condition was effective In addition to investigating the blinding for participants stimulation condition, we also conducted the same analyses for the pharmacological group allocations. These findings also supported our prediction that both participants and the experimenter would be unable to correctly identify whether they were in the levodopa or placebo condition. Overall, there was an bias for participants to report they received the placebo tablet. Specifically, 122 out of 231 participants correctly guessed their pharmacological condition (53%). In the levodopa condition, 34 out of 116 participants correctly guessed their group (29%) and 82 participants guessed incorrectly (70%). However, in the placebo condition 88 out of 115 participants correctly guessed their group (77%) and 27 participants guessed incorrectly (23%). This trend was in the opposite direction for the experimenter condition, whereby they were more accurate at classifying participants in the levodopa condition than in the placebo condition. Overall, the experimenter correctly classified 128 out of 231 into their pharmacological condition (55%). The experimenter classified 75 out of 116 participants correctly in the levodopa group (65%) and incorrectly classified 41 participants (35%). However, in the placebo group, the experimenter only correctly classified 53 out of 115 participants (46%) and incorrectly classified 62 participants (54%). Combined, these findings suggest that the pharmacological group blinding was effective for both the participants and the experimenter.

### Exploratory analyses

#### The effect of HD-tDCS on deliberately and automatically constrained thought

To assess the effect of HD-tDCS on deliberately and automatically constrained thought, we ran the same hierarchical order probit modelling as the central analyses above and followed up with assessing the stimulation block alone when there were discrepancies in the winning model selected by the LOOIC and Pseudo-BMA approaches. We hypothesised there may be a difference in the effect of stimulation, relative to the sham, across the placebo groups, for deliberately or automatically constrained thoughts. However, there was no evidence for an effect of HD-tDCS on deliberately or automatically constrained thought. The full output from these analyses can be found in the supplementary results: the top LOOIC and Pseudo-BMA model selection weights for the complete datasets and stimulation block datasets are in Table S5 and S6, respectively, and the model comparisons are in Figure S3. The winning predictors for each preferred LOOIC model are in Figure S4 and Table S7. The Bayesian independent samples t-tests results can also be found in Table S8.

#### The effect of dopamine on deliberately and automatically constrained thought

In addition to investigating the effects of HD-tDCS, we predicted there may be a difference in the effects of dopamine, relative to the placebo conditions for deliberately or automatically constrained thoughts. There was no evidence for levodopa affecting deliberately or automatically constrained thought.

#### The interactive effects of dopamine and stimulation on deliberately and automatically constrained thought

We predicted that the interaction between dopamine and stimulation may affect the reporting of deliberately or automatically constrained thoughts. For deliberately constrained thought, there was no evidence of an interactive effect of levodopa and HD-tDCS. There was also limited evidence to support any interactive effect of stimulation and dopamine on automatically constrained thought; there were no stimulation-dopamine effects selected for the complete dataset. In the stimulation block alone, there was a meaningful positive main effect of the stimulation-dopamine predictor (*b* = .49, 95% CI [.05, .93]), which suggests that when active stimulation was combined with levodopa, participants were more likely to have automatically constrained thoughts, compared to when participants received the placebo drug with active stimulation. Given the investigation into baseline differences for this thought type met our threshold to be considered meaningful and found moderate evidence for no group differences (BF_excl_ = 8.11), the stimulation block finding does provide initial evidence for an effect of levodopa and stimulation on automatically constrained thought. However, the t-test aligned with the complete dataset analyses, finding anecdotal evidence for a null effect (BF_01_ = 2.23). This effect did not meet our threshold to be considered meaningful, however, the combined investigation suggests there is only weak preliminary evidence for an interactive effect of stimulation and dopamine on automatically constrained thought.

#### Assessing the role of motivation

There is evidence that motivation levels can impact task performance outcomes (Brosowsky et al., 2020) and that dopamine levels are linked to motivation (Mohebi et al., 2019), however no analyses were specified for this variable at Stage 1. Thus, we conducted a post-hoc exploratory Bayesian between-subjects ANOVA and a Bayesian independent samples t-test. The ANOVA included motivation level as the dependent measure and the four groups as the independent variable (active or sham stimulation combined with either the levodopa or placebo drug). There strong evidence for excluding the group variable (BF_excl_ > 14.22) which indicates there were no differences in motivation levels between the four experimental groups. We also ran a Bayesian independent samples t-test which directly compared the levodopa and placebo conditions and included motivation as the dependent variable. This test also found moderate, meaningful evidence against any differences in motivation between the two groups (BF_01_ = 6.06). Together, these findings suggest motivation did not influence the current results, as it did not differ between the experimental groups.

#### Exploratory blinding assessment

To assess whether the blinding was meaningful for both the participants and the experimenter, we followed up from the analyses above by running an exploratory Bayesian independent samples t-test for each condition (stimulation and drug) and group (participant and experimenter). This test included the objective group classification as the independent variable and the participant or experimenter’s subjective classification as the respective dependent variable. For the participants stimulation condition, there was meaningful evidence for a null effect (BF_01_ = 6.41), which suggests participants were unable to distinguish their stimulation condition. For the experimenter, the t-test showed anecdotal evidence for an effect (BF_10_ = 2.37), however as this did not meet our threshold to be considered meaningful, there was inconclusive evidence for the experimenter blinding. There was also inconclusive evidence for the participant and experimenter blinding of the drug condition, with both tests finding anecdotal evidence against an effect (BF_01_ = 2.87 and BF_01_ = 1.89, respectively). Overall, these results suggest the blinding was successful as there were no meaningful differences in the participant or experimenter classification of the stimulation or drug conditions.

## Discussion

This study investigated the causal role of the left PFC and dopamine in mind wandering and internal thought processes using application of HD-tDCS in conjunction with a pharmacological manipulation. This was a double sham- and placebo-blinded study, designed to replicate the HD-tDCS effect found by Rasmussen et al. (2024) and to expand on this research to explore the unique and interactive role of the dopaminergic system in dynamic internal thoughts. We found no evidence for replicating the effect of 2mA anodal HD-tDCS reducing freely moving thought, nor an effect of stimulation on task unrelated thought. However, we did find evidence that levodopa may reduce freely moving thought. This finding provides insight into the role of the dopaminergic system in internal thought processes and aligns with previous research to suggest this measure of mind wandering is independent of task unrelated thought (Christoff et al., 2016; Kam et al., 2021; Mills et al., 2018) and may be uniquely affected by experimental manipulations investigating periods of internal thought (Kam et al., 2021; Rasmussen et al., 2024). We also found preliminary evidence for an opposing effect of stimulation alone, relative to active stimulation combined with levodopa on participants behavioural variability during periods of task unrelated thought. These behavioural findings indicate that dopamine may improve behavioural performance, when combined with stimulation, and they emphasise the importance of using more sensitive behavioural measures to be able to detect periods of internal thought.

The evidence against an effect of anodal stimulation on freely moving thought does not support our hypothesised effect of a reduction in this thought type with active stimulation, and this also fails to replicate the preliminary findings from Rasmussen et al. (2024). Further, this is now the third registered report study to fail to replicate the original finding by Boayue et al. (2021) of 2mA anodal stimulation reducing task unrelated thought (Alexandersen et al., 2022; Rasmussen et al., 2024). The failure to replicate previous findings highlights the importance of pre-registering studies and the necessity to replicate effects within the field of mind wandering to establish reliability. It may be that 2mA HD-tDCS does not modulate task unrelated thoughts. However, it may also be that the overarching measure of task unrelated thought is not sufficient to capture the unique effects of stimulation on the diverse range of internal thought processes, and thus a more nuanced exploration into distinct types of internal thought is needed to effectively target aspects of internal thought. This reinforces the importance of approaching mind wandering from a more heterogenous perspective to investigate and target the current direction of one’s thoughts (Kam et al., 2021; Martel et al., 2019).

The failure to find direct effects of stimulation to single cortical regions on freely moving or task unrelated thoughts may reflect the recruitment of broader brain networks for these processes, which are relatively robust to focal non-invasive brain stimulation. This would align with imaging evidence finding that both the frontoparietal control network (Christoff et al., 2009; Fox et al., 2015) and default mode network (Christoff et al., 2009; Fox et al., 2015; Groot et al., 2021) are implicated in mind wandering, as these networks communicate across brain regions. Indeed, there is recent evidence to suggest that tDCS is unable to modulate effective connectivity within and between the default mode network during periods of mind wandering (Coulborn & Fernández-Espejo, 2022). This research found periods of mind wandering were associated with increases in connectivity between the DLPFC and posterior cingulate cortex (PCC), alongside overall excitatory coupling between regions with each targeted brain network (Coulborn & Fernández-Espejo, 2022). Furthermore, there is evidence from EEG research that synchronization between brain regions associated with the default mode network (Kirschner et al., 2012), alongside synchronization between theta and alpha oscillations (Rodriguez-Larios & Alaerts, 2021) facilitate periods of mind wandering. Thus, while the current findings suggest there is no effect of HD-tDCS on distinct internal thought types, future research could investigate whether it is possible to modulate mind wandering by applying non-invasive brain stimulation techniques across brain networks. Alternatively, other methods such as cross-frequency tACS may be more effective at targeting mind wandering processes, as this method is able to affect the synchronisation between brain regions (Riddle et al., 2021; Turi et al., 2020).

This study adds to the inconsistences found across the tDCS and mind wandering literature, and thus it is also critical to consider the methodological practices that should be employed in this field. Firstly, the importance of replicating effects has been demonstrated, given both the current study and Alexandersen et al. (2022) failed to replicate previous findings. The original registered report by Rasmussen et al. (2024) and pre-registered study by Boayue et al. (2021) were arguably methodologically advanced studies, being high-powered studies with robust analytical approaches. However, the current study included a greater sample size (60 participants per group, instead of the 40 included in the previous study) and the probit modelling analyses were also completely established at Stage 1 acceptance, whereas it was included in the exploratory section of the previous paper, due to a lack of specificity in the interpretation of effects at Stage 1. This also highlights the importance of employing open sciences practices, including pre-registering studies or developing registered reports to improve confidence in scientific outcomes. In addition to replicating results, it is important to double-blind the participants and experimenter to the experimental conditions, to avoid any possibility of placebo effects driving the effects, and this was achieved successfully in the current study. Thus, to contribute meaningfully to the field of mind wandering using tDCS, it is critical to employ rigorous methodologies with replication studies being an essential component of the literature to improve confidence in empirical outcomes.

In this study, we were also interested in understanding the role of the dopaminergic system in mind wandering and dynamic thought, given there is evidence to link dopaminergic functioning to mind wandering (O’Callaghan et al., 2021) and attentional control processes (Cools, 2016; Cools & D’Esposito, 2011; Leow et al., 2024). Our hypothesis that there would be a difference in freely moving thought between the levodopa and placebo groups, across the sham condition, was supported. Specifically, there was evidence for levodopa reducing freely moving thought, relative to the placebo condition. It is important to note that this finding was established by a main effect of levodopa in the complete dataset, rather than the block x levodopa interaction, which means the data included the baseline block. However, this finding was also established as a robust effect in the stimulation block alone. Furthermore, it is unlikely that group differences in baseline performance could explain the main effect found for the complete dataset, as the groups were demographically balanced with large sample sizes (i.e. 60 participants in each group) and there was no evidence for any baseline group differences. The pharmacological drug blinding was also effective for both the participants and the experimenter, which suggests these results were not driven by a placebo effect. The finding supports preliminary imaging literature finding a relationship between the dopaminergic system and mind wandering (O’Callaghan et al., 2021). The dopaminergic system has been implicated in cognitive control processes (Cools, 2008, 2016); one possibly mechanism for this could be through facilitating sustained attention via reduced shifts towards internal thought. While there was a meaningful effect for freely moving thought, there was no evidence for an effect of levodopa on task unrelated thought. This suggests targeting the dopaminergic system may specifically affect more creative and spontaneous internal thought processes (Christoff et al., 2016), as freely moving thought is also independent of task relatedness (Kam et al., 2021; Mills et al., 2018; Rasmussen et al., 2024). Overall, these findings suggest that the dopaminergic system is implicated in shifts towards freely moving thought and they provide preliminary evidence that targeting this system may reduce periods of spontaneous, internal thought, during a cognitively demanding task.

Another key investigation in this study was understanding the interactive effects of levodopa and HD-tDCS on freely moving thought and task unrelated thought, as we predicted that this combination may facilitate or inhibit internal thought process. However, we did not find any evidence to support an interactive effect of dopamine and stimulation on either thought type. To date, there has been no investigation into this interaction in relation to mind wandering, as studies have primarily focused on the combined effects of dopamine and tDCS on motor learning (Curtin et al., 2024; Fresnoza et al., 2014) and decision making (Leow, Marcos, et al., 2023). While these results suggest there may not be a direct interactive effect on self-reported mind wandering, the current study did find an opposing effect of stimulation alone, compared to the interactive effects of stimulation and dopamine on participants behavioural variability during periods of task unrelated thought. Specifically, for HD-tDCS with placebo, those with active stimulation had greater variability in their responses during periods of mind wandering, compared to those who received sham stimulation. In contrast, for HD-tDCS with levodopa, those with active stimulation had less variability in their responses during periods of mind wandering. This aligns with research in other fields where the effect of tDCS is reversed with the administration of levodopa (e.g., motor sequence learning; Leow, Jiang, et al., 2023). Thus, whilst there was no evidence for interactive effects between stimulation and dopamine on propensity to mind wander, there was preliminary evidence that the behavioural consequences of mind wandering may show such an interaction.

The FT-RSGT has been proposed as a more sensitive measure of task performance, relative to other measures used in mind wandering research such as a SART (Alexandersen et al., 2022; Boayue et al., 2021). Indeed, regardless of stimulation the FT-RSGT showed sensitivity to periods of different types of internal thought as reflected in the opposing relationship between behavioural variability for deliberately constrained thought, compared to task unrelated and freely moving thought. Specifically, when participants were engaging in deliberately constrained thought, they achieved lower levels of variability (i.e. better performance). In contrast, for task unrelated and freely moving thought, variability was higher (i.e. performance was worse). Furthermore, greater variability was more predictive of task unrelated and freely moving thought when participants randomness was increased. However, the opposite interaction was true for deliberately constrained thought, such that when randomness increased, reduced behavioural variability was more predictive of these goal-directed thoughts. Overall, our findings suggest the FT-RSGT can discriminate between heterogeneous dynamic thought types on task performance during a cognitively demanding task. However, despite this apparent sensitivity it is important to note that these effects may change depending on the demands of the task, as there is evidence that individuals report more deliberate mind wandering during easy tasks and more spontaneous, unintentional mind wandering during difficult tasks (Seli, Konishi, et al., 2018). The influence of task difficulty on the propensity and nature of mind wandering performance – alongside the potential for this factor to affect HD-tDCS and levodopa manipulations – would be an interesting route of investigation for future studies in this field.

In summary, this study found no evidence for a direct effect of left PFC HD-tDCS on modulating freely moving, nor task unrelated thought. However, there was preliminary evidence that levodopa may reduce freely moving thought, which supports the role of the dopaminergic system in internal thought processes. Furthermore, this research found that dopamine may moderate the relationship between stimulation and behavioural variability performance during periods of task unrelated thought, which supports the interactive role of the dopaminergic system in affecting behavioural tDCS outcomes. These findings highlighted the importance of employing sensitive behavioural tasks, such as the FT-RSGT, to detect periods of internal thought, and this may be able to overcome some of the limitations associated with self-reported measures of mind wandering. The failure to replicate previous HD-tDCS findings further emphasises the importance of employing robust methodological practices to improve confidence in the results within this field. It is also important to consider alternative avenues which could be used to affect mind wandering, such as targeting multiple brain regions or employing alternative non-invasive brain stimulation methods. Finally, the dopaminergic findings provide preliminary evidence for the potential effectiveness of targeting this system to reduce internal thoughts and improve behavioural performance, which may be able to be used in contexts which require sustained attention.

## Supporting information

Supplementary Results

## Footnotes

1 In line with previous research using dopamine manipulations in our lab (Leow, Jiang, et al., 2023; Leow, Marcos, et al., 2023), the placebo tablet used was a vitamin C tablet, instead of a multivitamin, due to the closer colour similarity to the levodopa tablet.

2 There was one participant who had their blood pressured measured pre-drug administration and 35 minutes after administration, but the final measure was missed. This participant was removed from the blood pressure and heart rate analyses; however, they were included in all other analyses as they met the ethical requirements (for recording blood pressure measures) to be included in the study. Furthermore, there was one additional participant whose response to the initial BL-VAS failed to record correctly, and thus they were excluded from the mood analysis, but included in all other analyses for the study.

