## Supplementary Results for "On the neural substrates of mind wandering and dynamic thought: A drug and brain stimulation study"

### Figure S1. Model comparisons for each set of probit models

A. Freely Moving Thought – Active vs Sham (across Placebo conditions)
*Complete Dataset Winning Models (LOOIC and Pseudo-BMA methods agreed)*

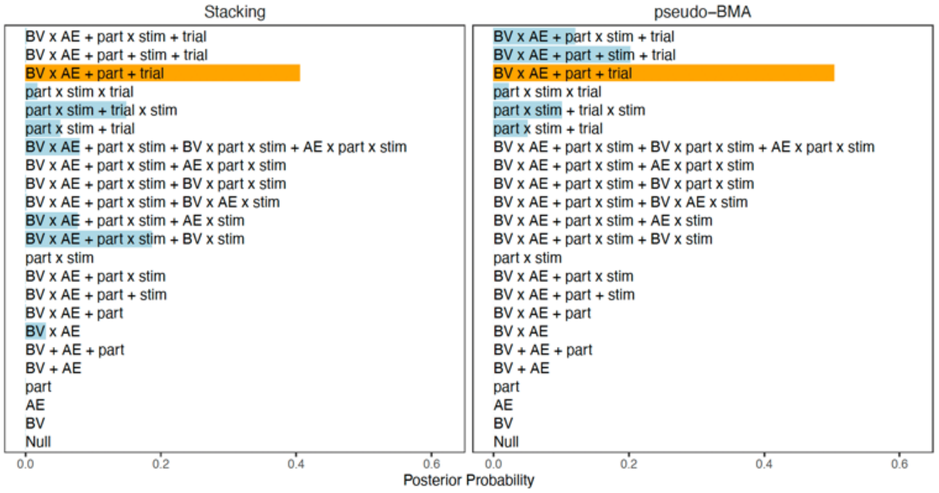


B. Task Unrelated Thought – Active vs Sham (across Placebo conditions)
*Complete Dataset Winning Models (LOOIC and Pseudo-BMA methods disagreed)*

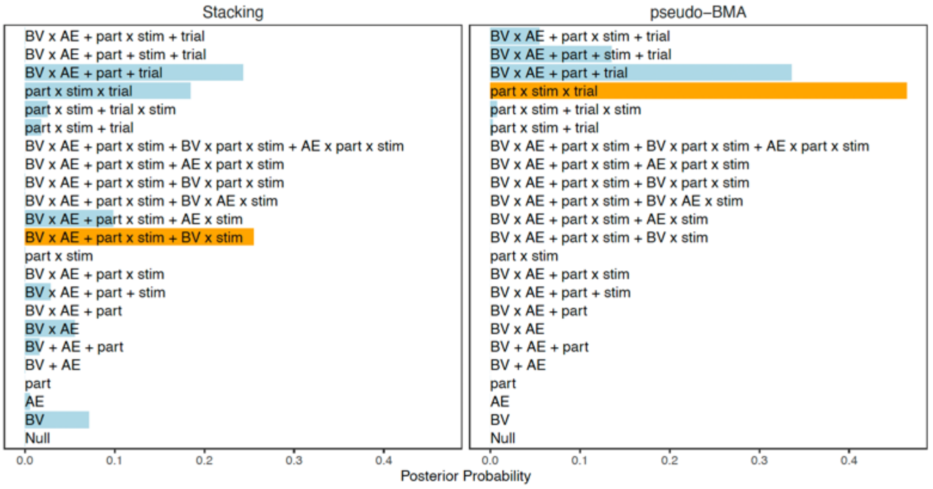


*Stimulation Dataset Winning Models (LOOIC and Pseudo-BMA methods agreed)*

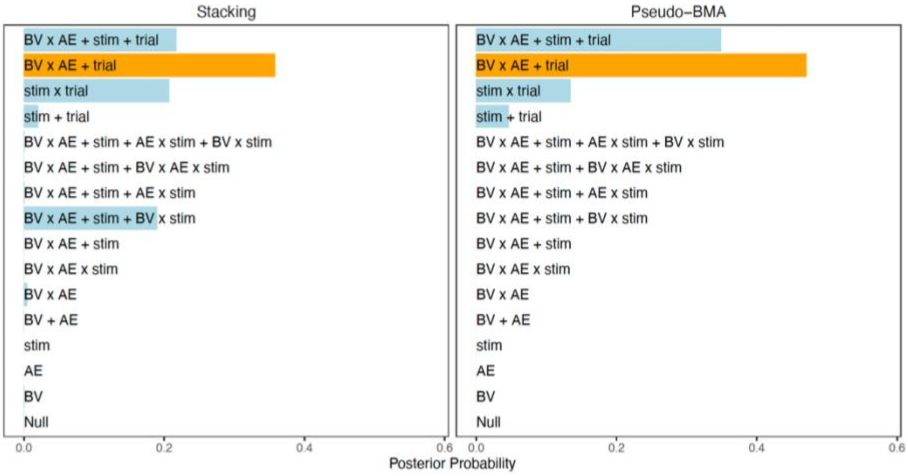

C. Freely Moving Thought – Levodopa vs Placebo (across Sham conditions)

*Complete Dataset Winning Models (LOOIC and Pseudo-BMA methods disagreed)*

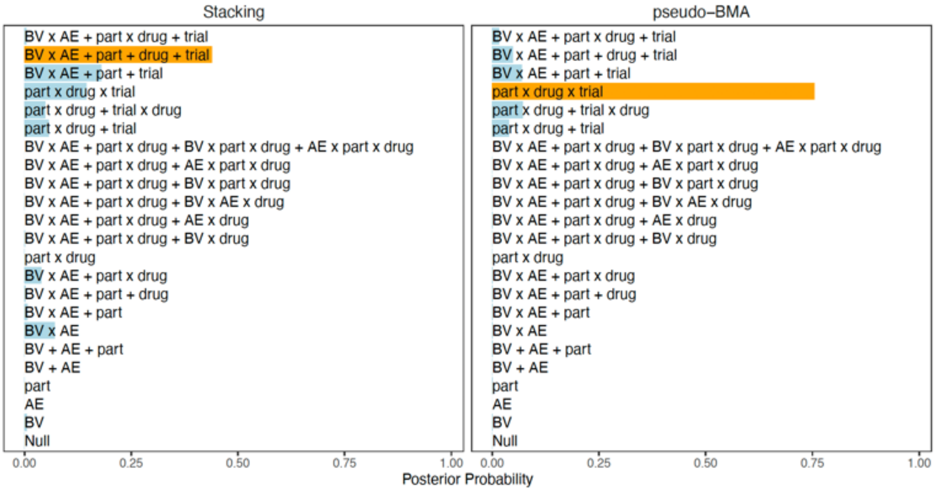


*Stimulation Dataset Winning Models (LOOIC and Pseudo-BMA methods disagreed)*

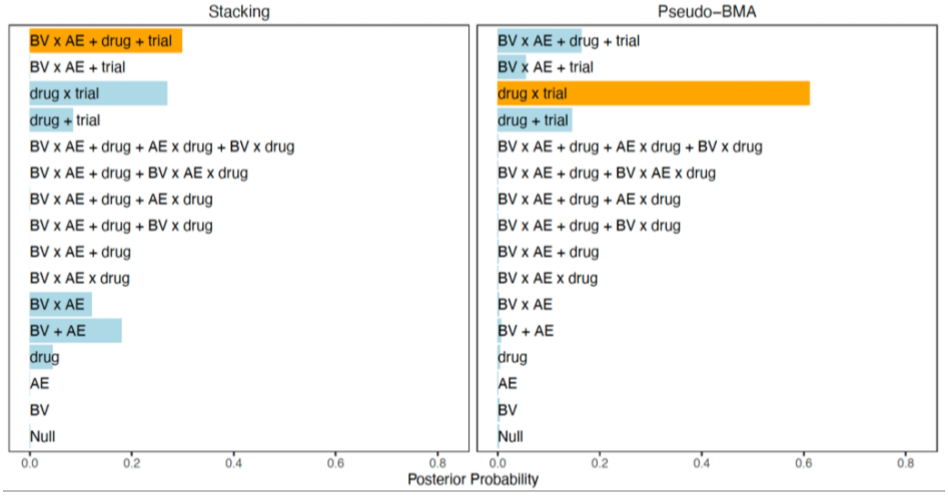


D. Task Unrelated Thought – Levodopa vs Placebo (across Sham conditions)

*Complete Dataset Winning Models (LOOIC and Pseudo-BMA methods disagreed)*

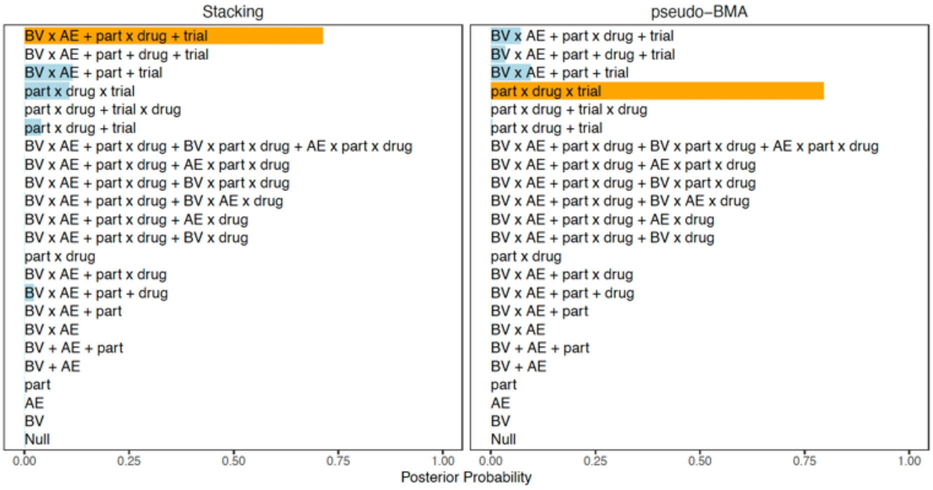

*Stimulation Dataset Winning Models (LOOIC and Pseudo-BMA methods agreed)*

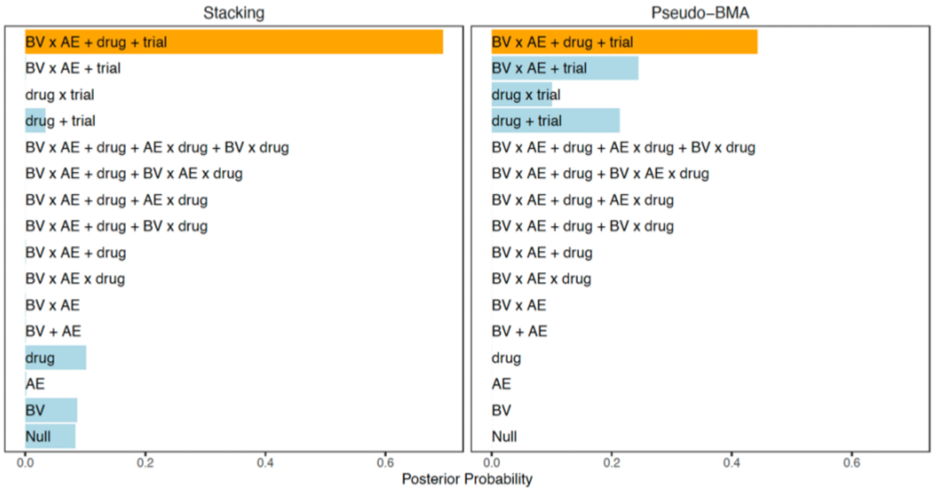


E. Freely Moving Thought – Active Stim with Levodopa vs Active Stim with Placebo
*Complete Dataset Winning Models (LOOIC and Pseudo-BMA methods agreed)*

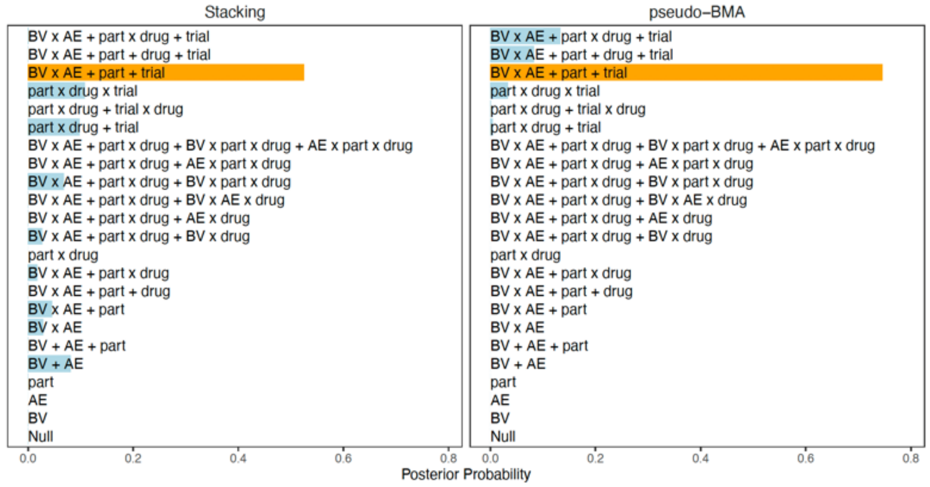


F. Task Unrelated Thought – Active Stim with Levodopa vs Active Stim with Placebo
*Complete Dataset Winning Models (LOOIC and Pseudo-BMA methods disagreed)*

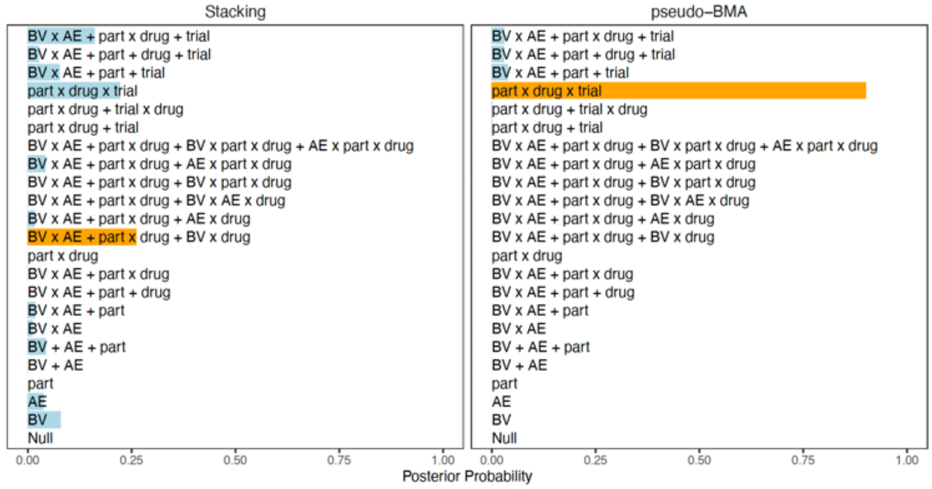

*Stimulation Dataset Winning Models (LOOIC and Pseudo-BMA methods agreed)*

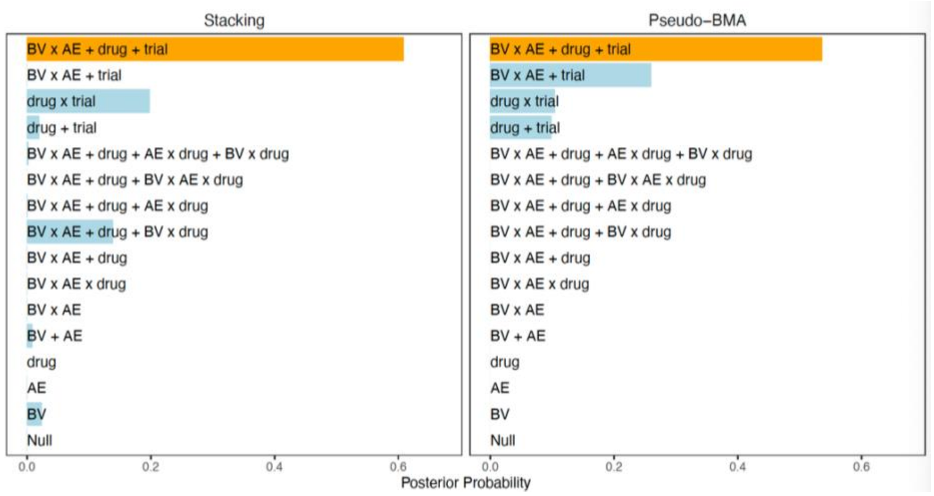


### Figure S2. Winning Model Predictors

A. Freely Moving Thought – Active vs Sham (across Placebo conditions)
*Complete Dataset Winning Models (LOOIC and Pseudo-BMA methods agreed)*

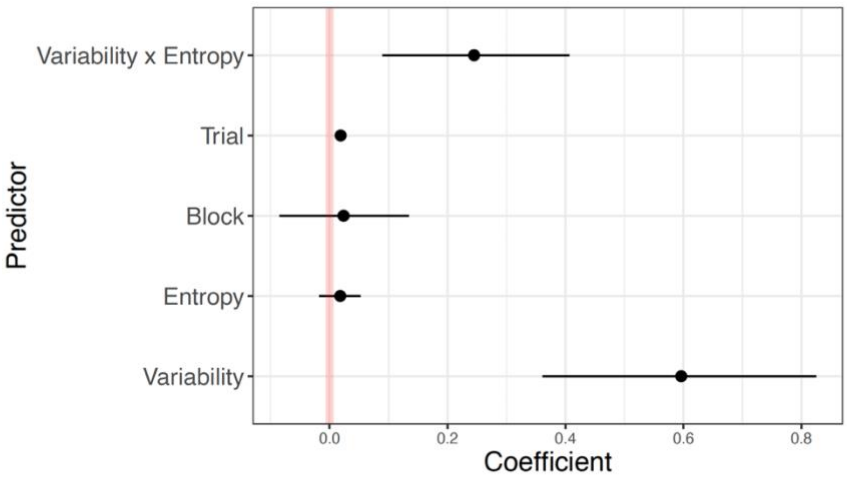


B. Task Unrelated Thought – Active vs Sham (across Placebo conditions)
*Complete Dataset Winning Models (LOOIC and Pseudo-BMA methods disagreed)*

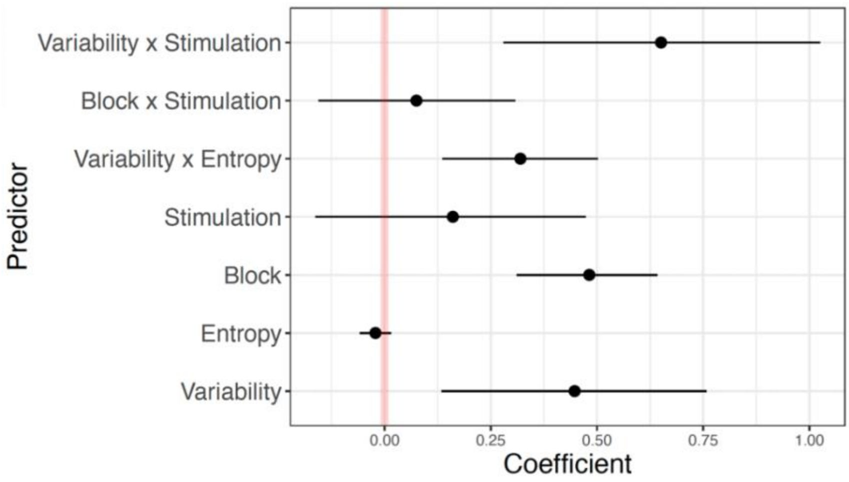

*Stimulation Dataset Winning Models (LOOIC and Pseudo-BMA methods agreed)*

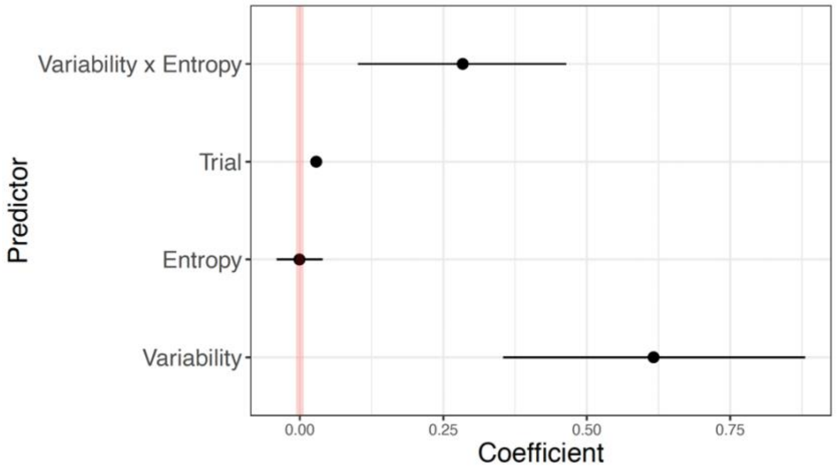


C. Freely Moving Thought – Levodopa vs Placebo (across Sham conditions)

*Complete Dataset Winning Models (LOOIC and Pseudo-BMA methods disagreed)*

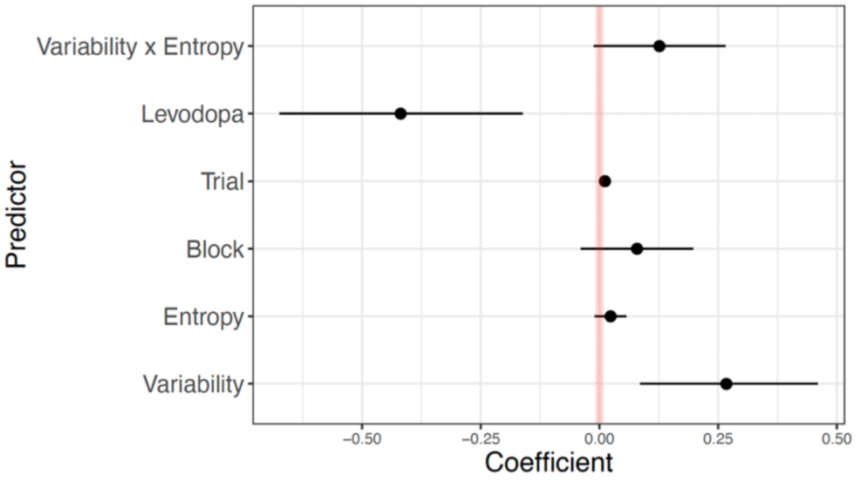


*Stimulation Dataset Winning Models (LOOIC and Pseudo-BMA methods disagreed)*

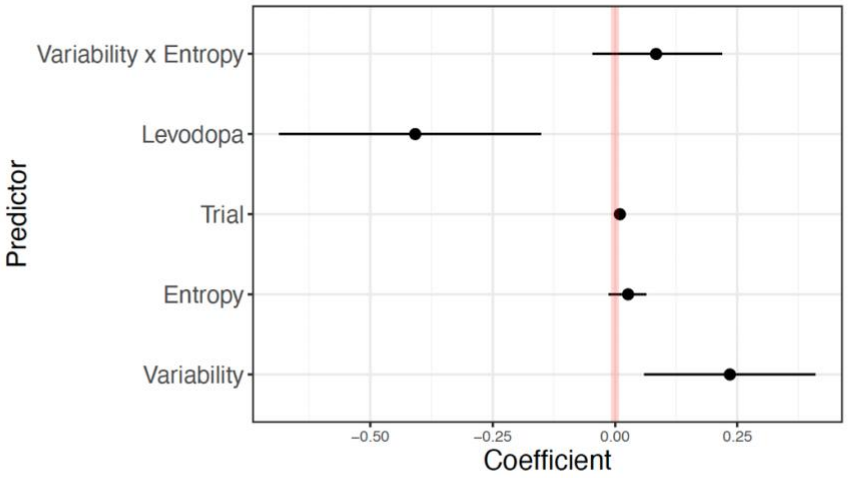

D. Task Unrelated Thought – Levodopa vs Placebo (across Sham conditions)

*Complete Dataset Winning Models (LOOIC and Pseudo-BMA methods disagreed)*

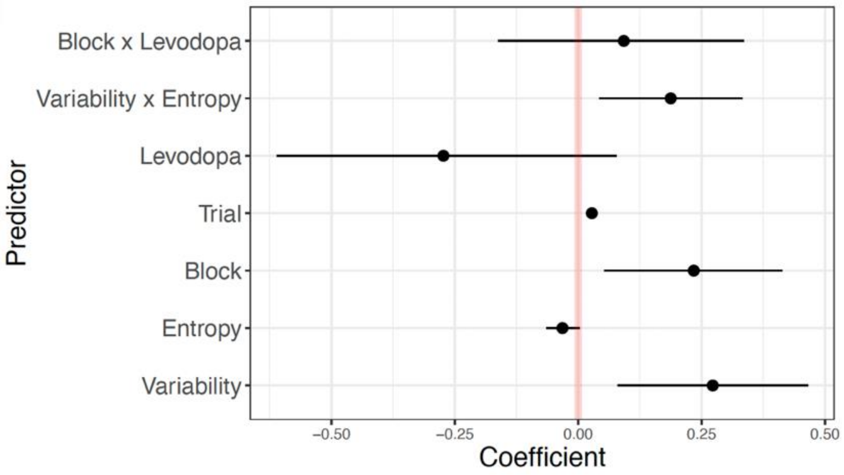


*Stimulation Dataset Winning Models (LOOIC and Pseudo-BMA methods agreed)*

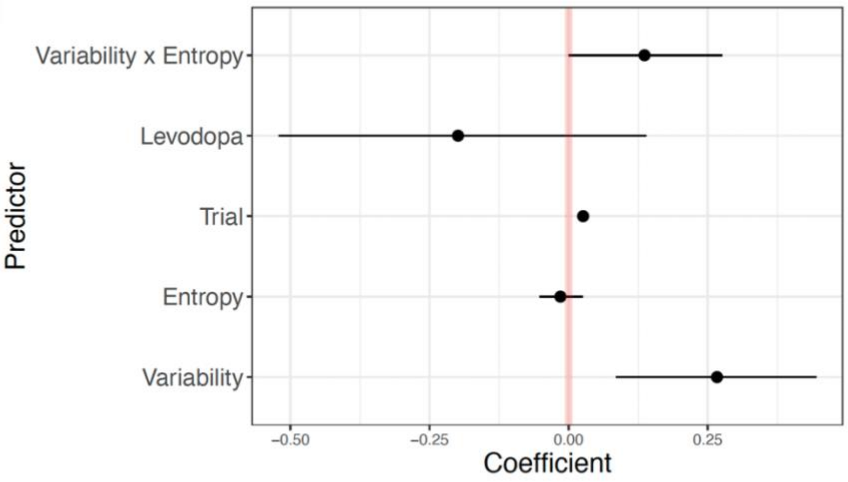


E. Freely Moving Thought – Active Stim with Levodopa vs Active Stim with Placebo

*Complete Dataset Winning Models (LOOIC and Pseudo-BMA methods agreed)*

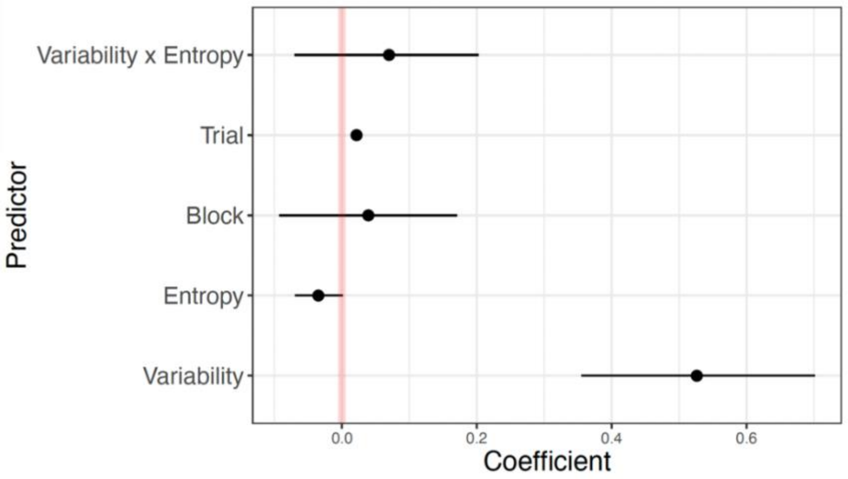

F. Task Unrelated Thought – Active Stim with Levodopa vs Active Stim with Placebo

*Complete Dataset Winning Models (LOOIC and Pseudo-BMA methods disagreed)*

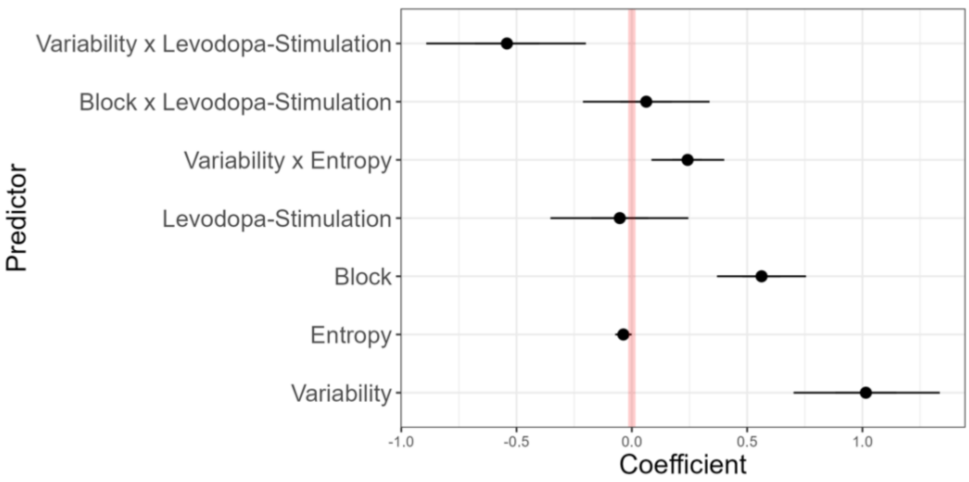


*Stimulation Dataset Winning Models (LOOIC and Pseudo-BMA methods disagreed)*

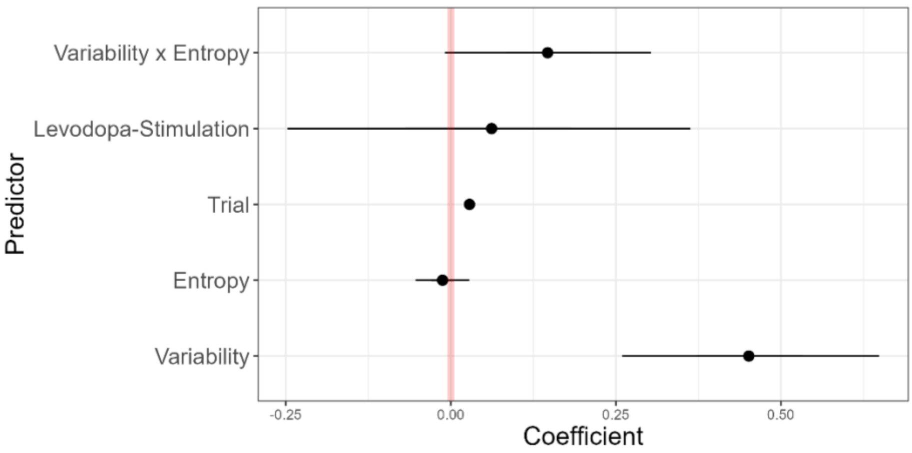


#### Table S1. Complete dataset selected model-coefficients

| Thought type | Active vs Sham Stimulation (Across Placebo Condition) | | Levodopa vs Placebo (Across Sham Condition) | | Active with Levodopa vs Active with Placebo | |
|---|---|---|---|---|---|---|
| | **LOOIC** | **Pseudo-BMA** | **LOOIC** | **Pseudo-BMA** | **LOOIC** | **Pseudo-BMA** |
| **Task unrelated thought** | Mod11 – 0.25 | Mod19 – 0.46 | Mod22 – 0.71 | Mod19 – 0.79 | Mod11 – 0.26 | Mod19 – 0.90 |
| | Mod20 – 0.24 | Mod20 – 0.34 | Mod20 – 0.11 | Mod20 – 0.10 | Mod19 – 0.22 | Mod20 – 0.04 |
| **Freely moving thought** | Mod20 – 0.41 | Mod20 – 0.50 | Mod21 – 0.44 | Mod19 – 0.75 | Mod20 – 0.52 | Mod20 – 0.75 |
| | Mod11 – 0.19 | Mod21 – 0.20 | Mod20 – 0.18 | Mod18 – 0.07 | Mod19 – 0.11 | Mod22 – 0.13 |

#### Table S2. Stimulation block dataset selected model-coefficients

| Thought type | Active vs Sham Stimulation (Across Placebo Condition) | | Levodopa vs Placebo (Across Sham Condition) | | Active with Levodopa vs Active with Placebo | |
|---|---|---|---|---|---|---|
| | LOOIC | Pseudo-BMA | LOOIC | Pseudo-BMA | LOOIC | Pseudo-BMA |
| Task unrelated thought | Mod14 – 0.36 | Mod14 – 0.47 | Mod15 – 0.70 | Mod15 – 0.44 | Mod15 – 0.61 | Mod15 – 0.54 |
| | Mod15 – 0.22 | Mod15 – 0.35 | Mod03 – 0.10 | Mod14 – 0.24 | Mod13 – 0.20 | Mod14 – 0.26 |
| Freely moving thought | | | Mod15 – 0.30 | Mod13 – 0.61 | | |
| | | | Mod13 – 0.27 | Mod15 – 0.16 | | |

#### Table S3. Winning LOOIC model from each set of probit models

A. Freely Moving Thought – Active vs Sham (across Placebo conditions)

**Complete Dataset**

| Predictor | Estimate | 95% Credible Intervals (CIs) |
|---|---|---|
| Behavioural variability | .59 | [.32, .87]* |
| Randomness | .02 | [-.03, .06] |
| Block | .02 | [-.10, .15] |
| Trial | .02 | [.01, .02] |
| Behavioural variability x Randomness | .25 | [.06, .44]* |

B. Task Unrelated Thought – Active vs Sham (across Placebo conditions)

**Complete Dataset**

| Predictor | Estimate | 95% CIs |
|---|---|---|
| Behavioural variability | .45 | [.08, .82]* |
| Randomness | -.02 | [-.07, .02] |
| Block | .48 | [.28, .68]* |
| Stimulation | .16 | [-.23, .53] |
| Behavioural variability x Randomness | .32 | [.11, .54]* |
| Block x Stimulation | .08 | [-.20, .35] |
| Behavioural variability x Stimulation | .65 | [.21, 1.09] |

**Stimulation Block Dataset**

| Predictor | Estimate | 95% CIs |
|---|---|---|
| Behavioural variability | .62 | [.31, .93]* |
| Randomness | -.00 | [-.05, .05] |
| Trial | .03 | [.02, .03]* |
| Behavioural variability x Randomness | .28 | [.07, .50]* |

C. Freely Moving Thought – Levodopa vs Placebo (across Sham conditions)

**Complete Dataset**

| Predictor | Estimate | 95% CIs |
|---|---|---|
| Behavioural variability | .27 | [.05, .49]* |
| Randomness | .02 | [-.02, .06] |
| Block | .08 | [-.06, .22] |
| Trial | .01 | [.01, .02]* |
| Behavioural variability x Randomness | .13 | [-.04, .29] |
| Levodopa | -.42 | [-.72, -.11]* |

**Stimulation Block Dataset**

| Predictor | Estimate | 95% CIs |
|---|---|---|
| Behavioural variability | .24 | [.03, .44]* |
| Randomness | .03 | [-.02, .07] |
| Trial | .01 | [.01, .01]* |
| Levodopa | -.41 | [-.74, -.10]* |
| Behavioural variability x Randomness | .08 | [-.07, .25] |

D. Task Unrelated Thought – Levodopa vs Placebo (across Sham conditions)

**Complete Dataset**

| Predictor | Estimate | 95% CIs |
|---|---|---|
| Behavioural variability | .27 | [.05, .50]* |
| Randomness | -.03 | [-.07, .01] |
| Block | .23 | [.02, .45]* |
| Trial | .03 | [.02, .03]* |
| Behavioural variability x Randomness | .19 | [.02, .36]* |
| Levodopa | -.27 | [-.68, .13] |
| Block x Levodopa | .09 | [-.21, .39] |

**Stimulation Block Dataset**

| Predictor | Estimate | 95% CIs |
|---|---|---|
| Behavioural variability | .27 | [.05. .48]* |
| Randomness | -.01 | [-.06, .03] |
| Trial | .03 | [.02, .03]* |
| Levodopa | -.20 | [-.57, .20] |
| Behavioural variability x Randomness | .14 | [-.03, .30] |

E. Freely Moving Thought – Active Stimulation with Levodopa vs with Placebo

**Complete Dataset**

| Predictor | Estimate | 95% Credible Intervals (CIs) |
|---|---|---|
| Behavioural variability | .53 | [.32, .73]* |
| Randomness | -.04 | [-.08, .01] |
| Block | .04 | [-.12, .19] |
| Trial | .02 | [.02, .03]* |
| Behavioural variability x Randomness | .07 | [-.10, .23]* |

F. Task Unrelated Thought – Active Stimulation with Levodopa vs with Placebo

**Complete Dataset**

| Predictor | Estimate | 95% CIs |
|---|---|---|
| Behavioural variability | 1.02 | [.64, 1.40]* |
| Randomness | -.04 | [-.08, .01] |
| Block | .56 | [.34, .79]* |
| Levodopa | -.05 | [-.42, .31] |
| Behavioural variability x Randomness | .24 | [.06, .43]* |
| Block x Levodopa | .06 | [-.26, .38] |
| Randomness x Levodopa | -.54 | [-.96, -.14]* |

**Stimulation Block Dataset**

| Predictor | Estimate | 95% CIs |
|---|---|---|
| Behavioural variability | .45 | [.22, .69]* |
| Randomness | -.01 | [-.06, .04] |
| Trial | .03 | [.02, .03]* |
| Levodopa | .06 | [-.30, .42] |
| Behavioural variability x Randomness | .15 | [-.04, .34] |

#### Table S4. Physiological measures pre- and post-drug administration

| | Levodopa | | | | Placebo | | | |
|---|---|---|---|---|---|---|---|---|
| | Pre | | Post | | Pre | | Post | |
| | *M* (SD) | *n* | *M* (SD) | *n* | *M* (SD) | *n* | *M* (SD) | *n* |
| **Mood** | 86.44 (5.85) | 115 | 88.70 (4.74) | 116 | 87.02 (6.08) | 115 | 88.32 (5.65) | 115 |
| **Systolic Blood Pressure** | 104.93 (13.01) | 116 | 100.28 (12.23) | 116 | 104.76 (14.19) | 115 | 103.25 (13.04) | 114 |
| **Diastolic Blood Pressure** | 73.97 (10.12) | 116 | 71.97 (9.12) | 116 | 74.45 (10.77) | 115 | 73.32 (9.45) | 114 |
| **Heart rate** | 79.99 (13.71) | 116 | 78.11 (12.95) | 116 | 79.70 (19.15) | 115 | 75.91 (15.68) | 114 |

#### Figure S3. Exploratory model comparisons for each set of probit models

A. Deliberately Constrained Thought – Active vs Sham (across Placebo conditions)
*Complete Dataset Winning Models (LOOIC and Pseudo-BMA methods disagreed)*

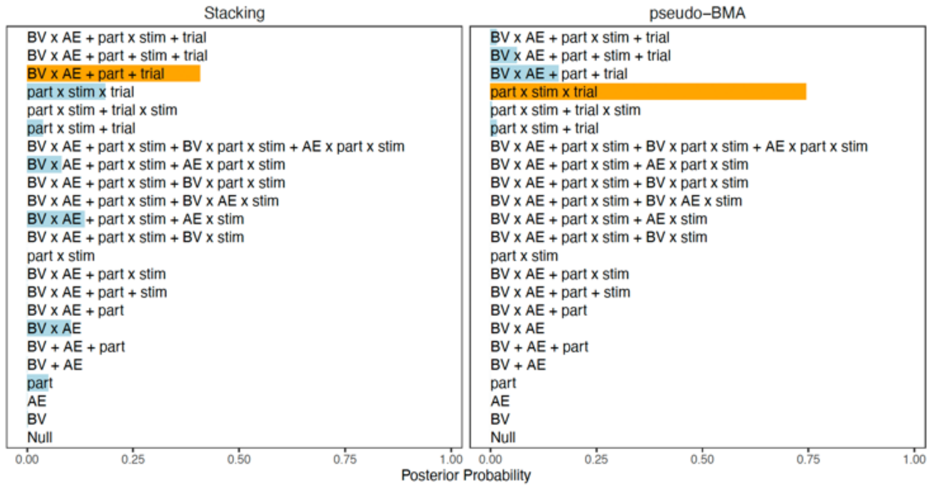


*Stimulation Dataset Winning Models (LOOIC and Pseudo-BMA methods disagreed)*

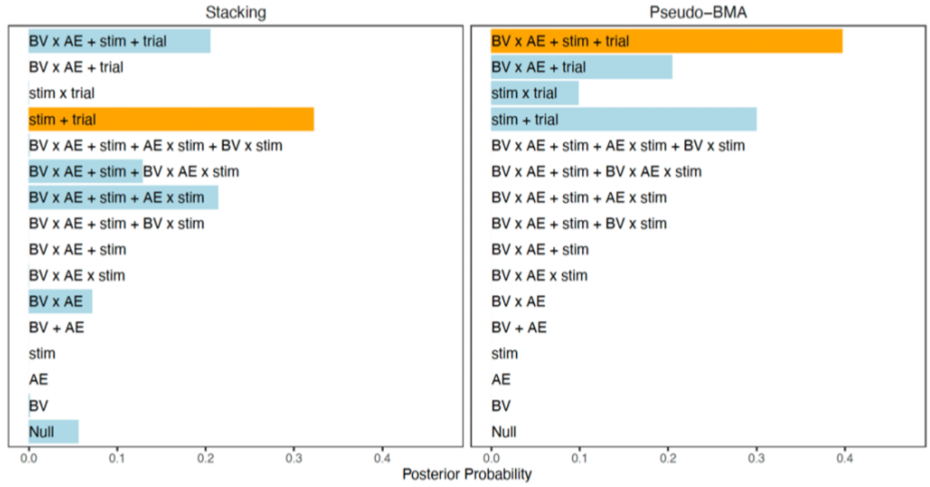

B. Automatically Constrained Thought – Active vs Sham (across Placebo conditions)

*Complete Dataset Winning Models (LOOIC and Pseudo-BMA methods disagreed)*

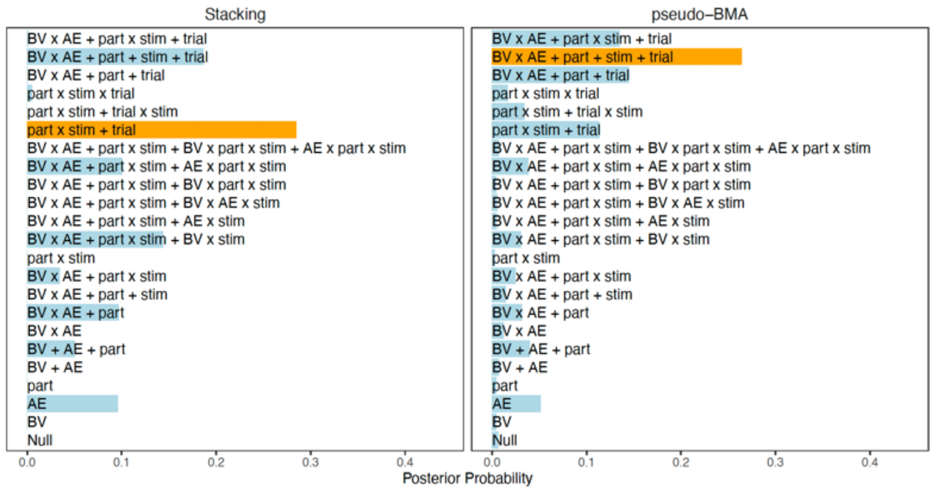


*Stimulation Dataset Winning Models (LOOIC and Pseudo-BMA methods agreed)*

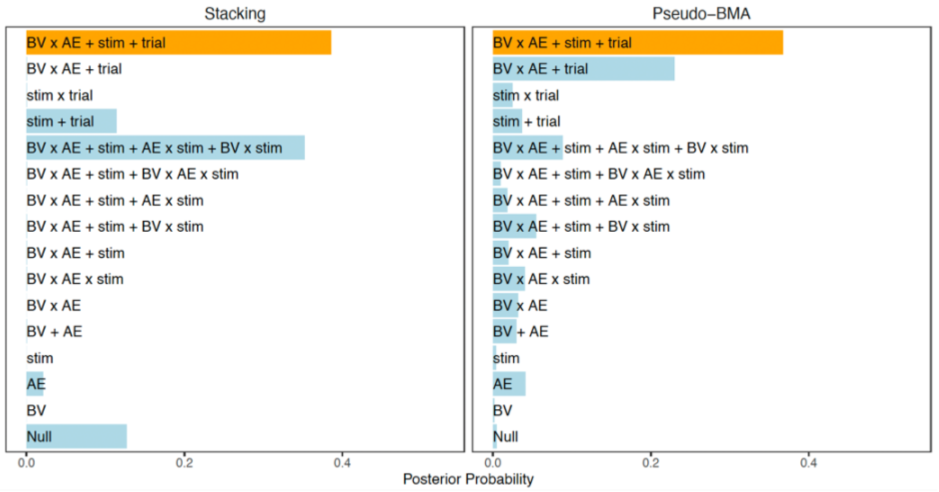


C. Deliberately Constrained Thought – Levodopa vs Placebo (across Sham conditions)

*Complete Dataset Winning Models (LOOIC and Pseudo-BMA methods disagreed)*

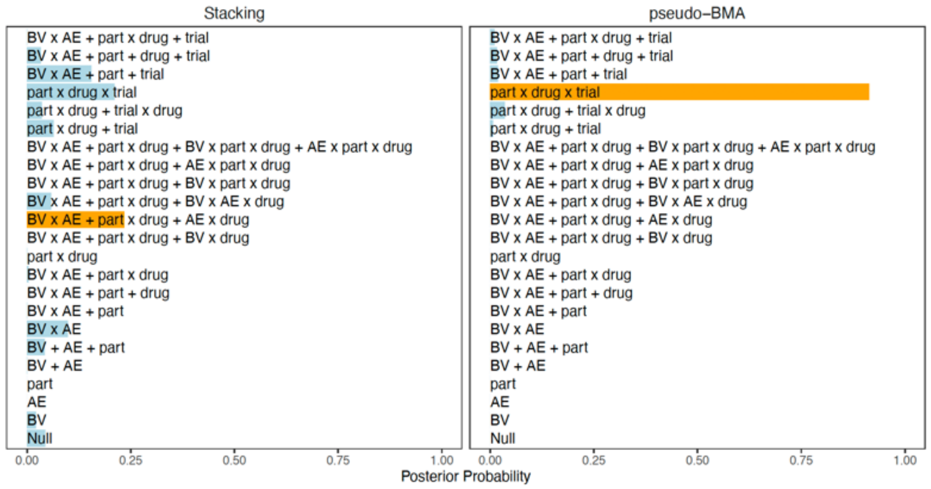

*Stimulation Dataset Winning Models (LOOIC and Pseudo-BMA methods agreed)*

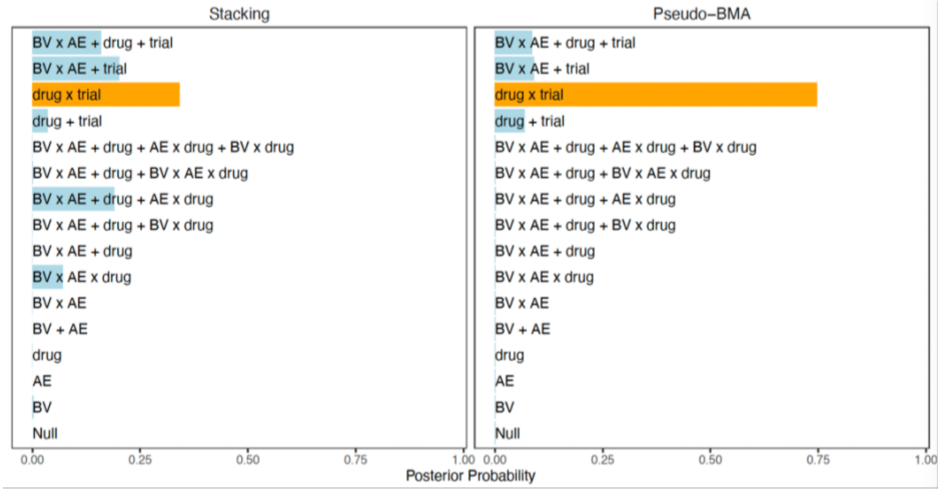


D. Automatically Constrained Thought – Levodopa vs Placebo (across Sham conditions)
*Complete Dataset Winning Models (LOOIC and Pseudo-BMA methods disagreed)*

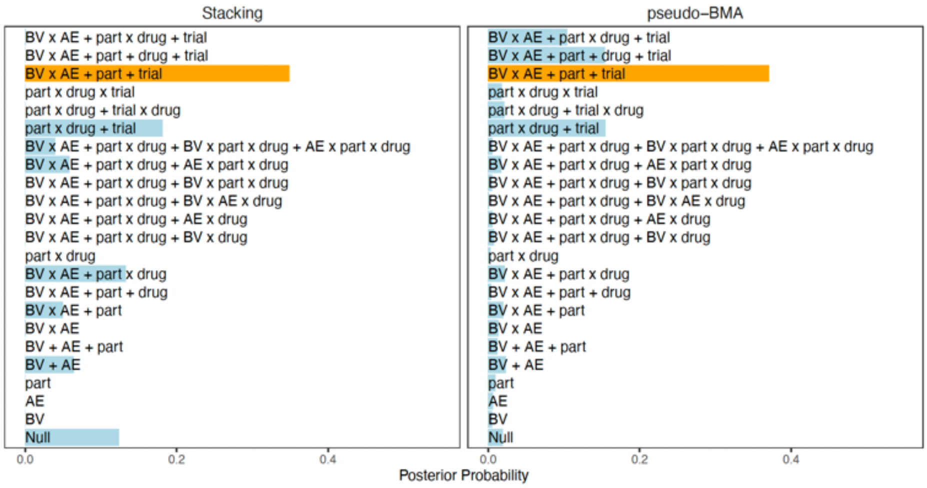


E. Deliberately Constrained Thought – Active Stim with Levodopa vs Active Stim with Placebo
*Complete Dataset Winning Models (LOOIC and Pseudo-BMA methods disagreed)*

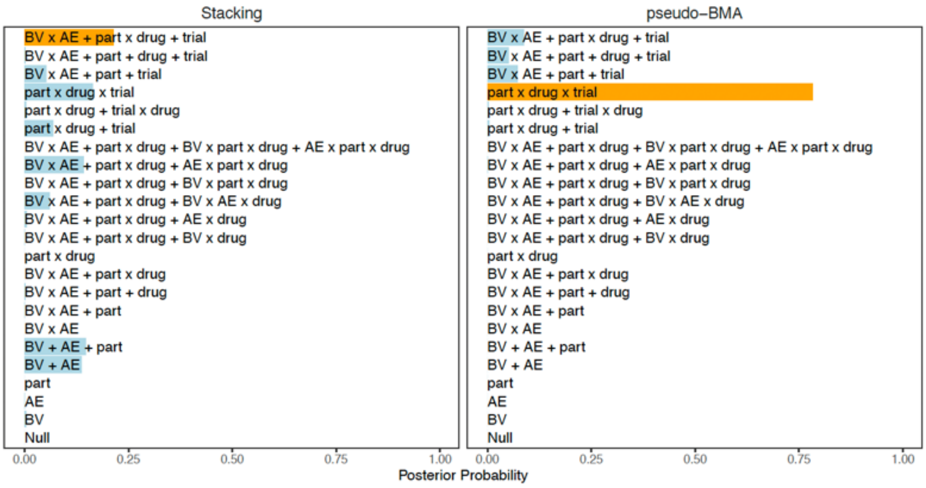

*Stimulation Dataset Winning Models (LOOIC and Pseudo-BMA methods disagreed)*

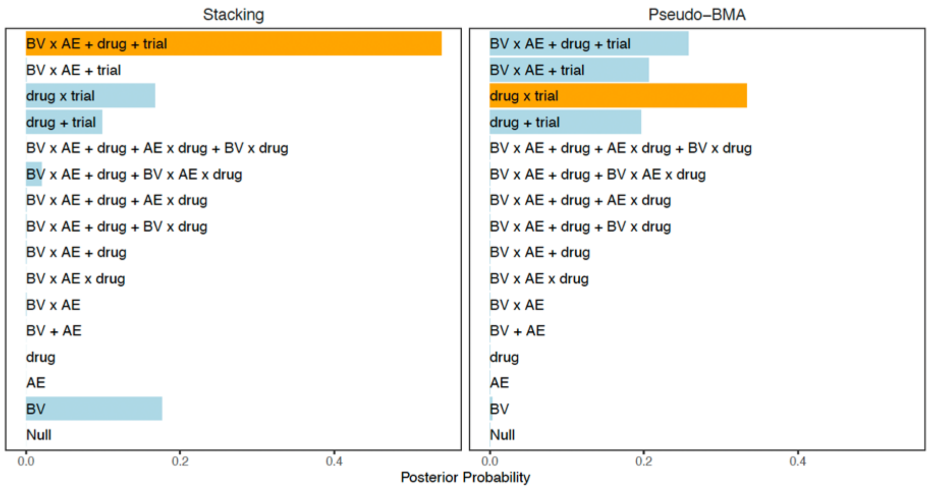


F. Automatically Constrained Thought – Active Stim with Levodopa vs Active Stim with Placebo

*Complete Dataset Winning Models (LOOIC and Pseudo-BMA methods disagreed)*

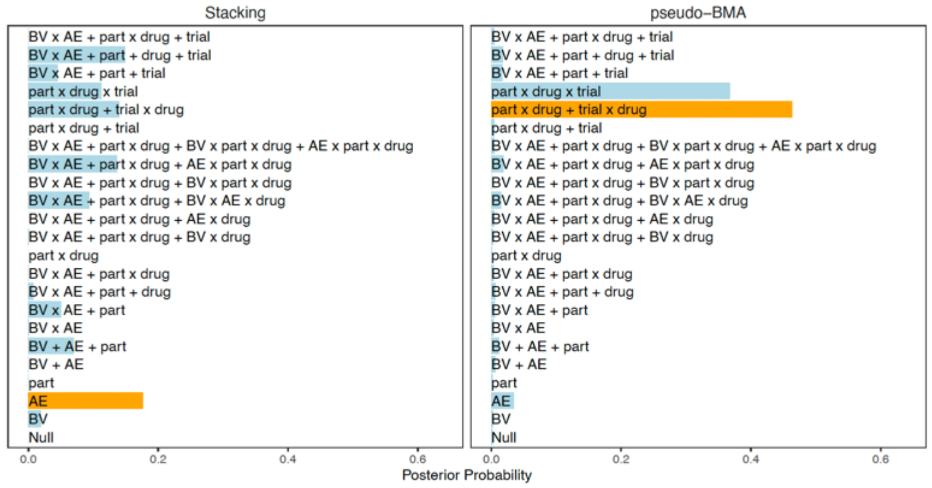


*Stimulation Dataset Winning Models (LOOIC and Pseudo-BMA methods agreed)*

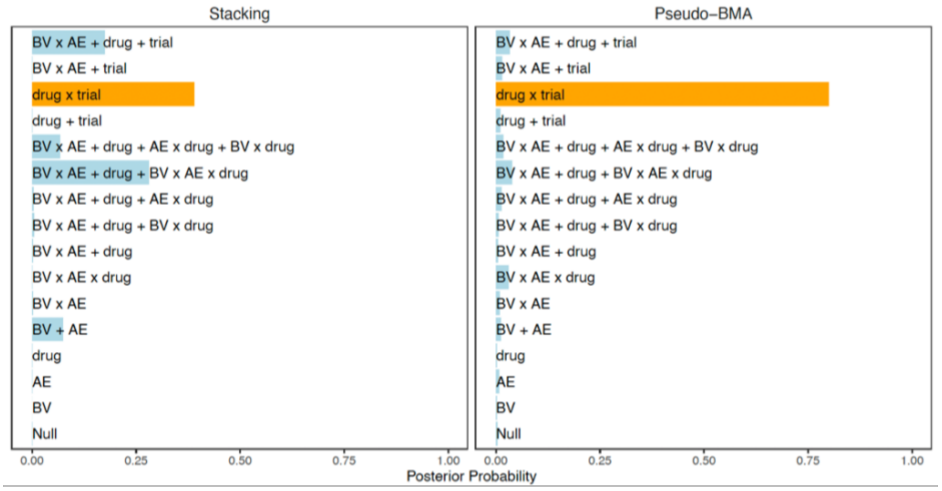

#### Figure S4. Exploratory Winning Model Predictors

A. Deliberately Constrained Thought – Active vs Sham (across Placebo conditions)

*Complete Dataset Winning Models (LOOIC and Pseudo-BMA methods disagreed)*

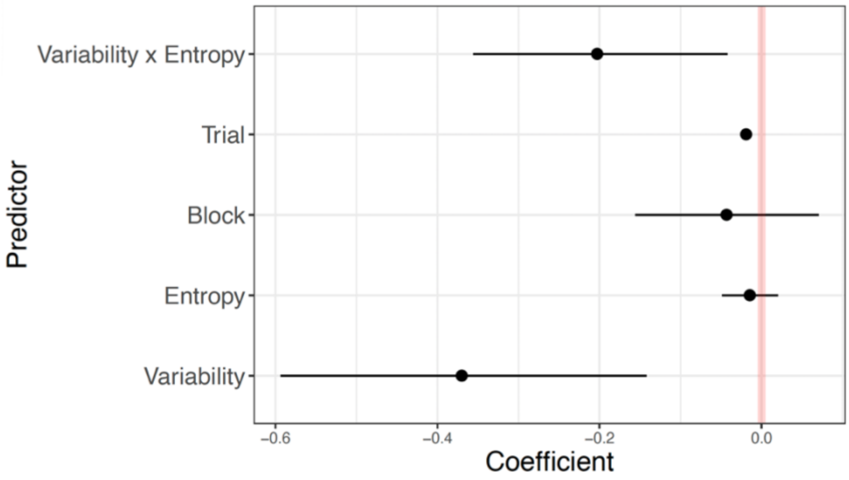


*Stimulation Dataset Winning Models (LOOIC and Pseudo-BMA methods disagreed)*

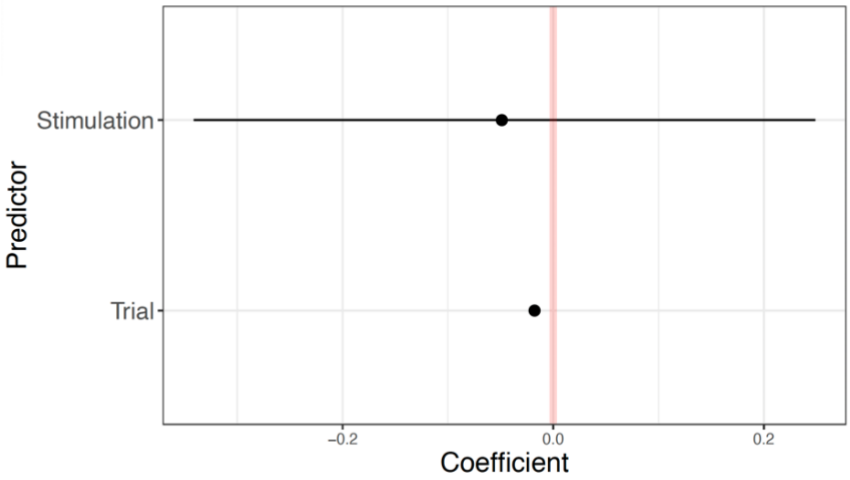


B. Automatically Constrained Thought – Active vs Sham (across Placebo conditions)

*Complete Dataset Winning Models (LOOIC and Pseudo-BMA methods disagreed)*

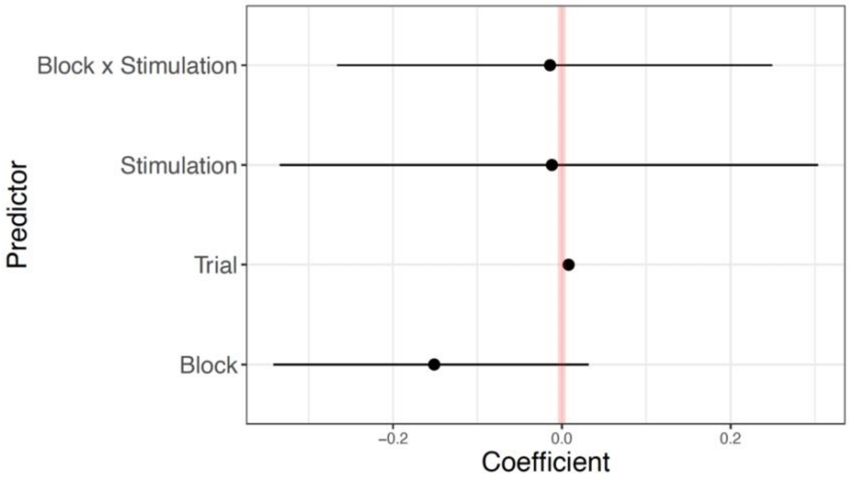

*Stimulation Dataset Winning Models (LOOIC and Pseudo-BMA methods agreed)*

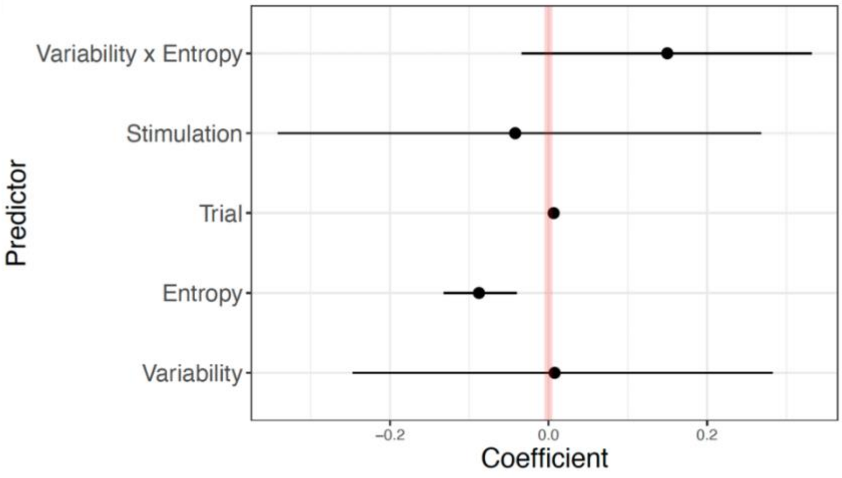


C. Deliberately Constrained Thought – Levodopa vs Placebo (across Sham conditions)
*Complete Dataset Winning Models (LOOIC and Pseudo-BMA methods disagreed)*

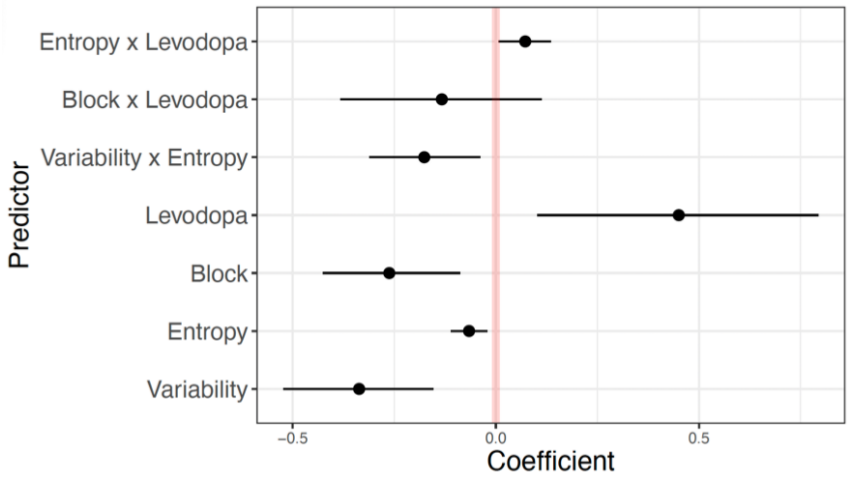


*Stimulation Dataset Winning Models (LOOIC and Pseudo-BMA methods agreed)*

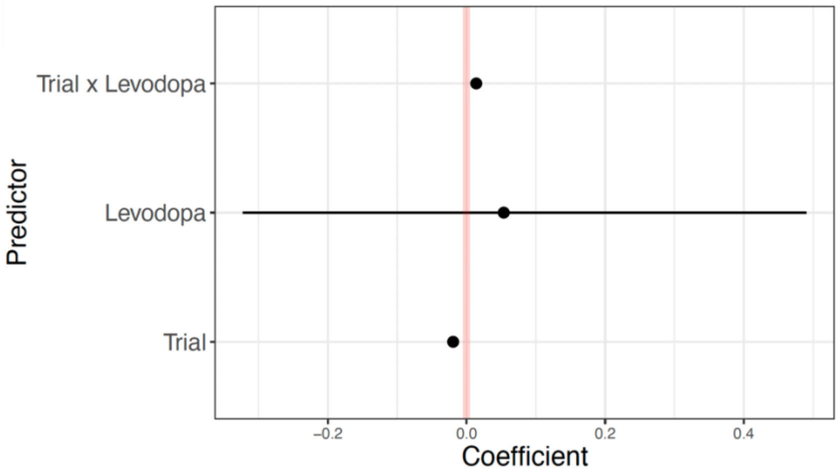

D. Automatically Constrained Thought – Levodopa vs Placebo (across Sham conditions)

*Complete Dataset Winning Models (LOOIC and Pseudo-BMA methods disagreed)*

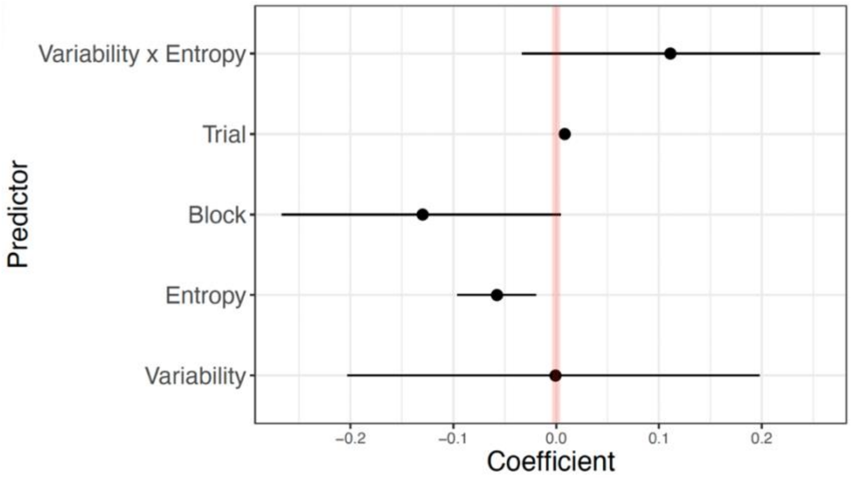


E. Deliberately Constrained Thought – Active Stim with Levodopa vs Active Stim with Placebo

*Complete Dataset Winning Models (LOOIC and Pseudo-BMA methods disagreed)*

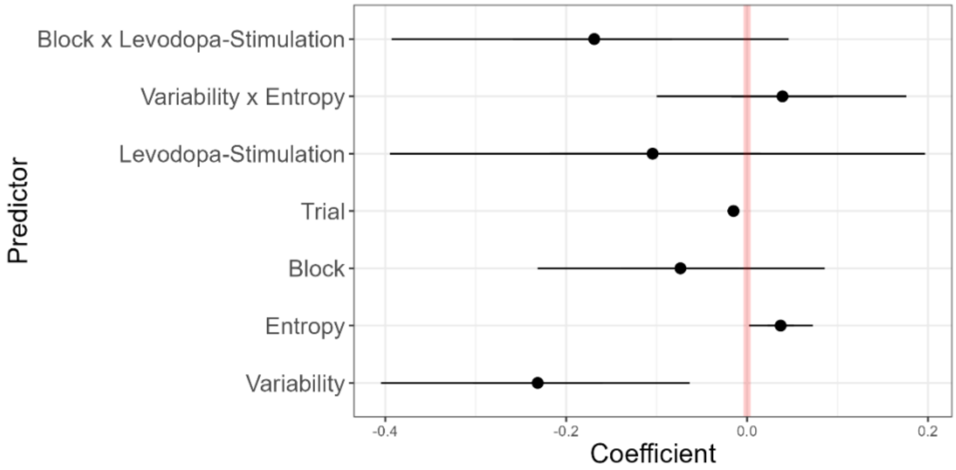


*Stimulation Dataset Winning Models (LOOIC and Pseudo-BMA methods agreed)*

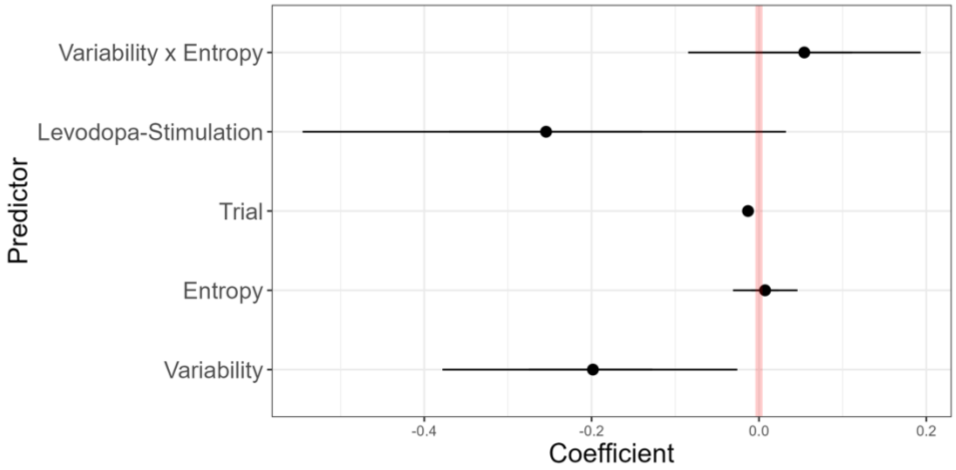

F. Automatically Constrained Thought – Active Stim with Levodopa vs Active Stim with Placebo

*Complete Dataset Winning Models (LOOIC and Pseudo-BMA methods disagreed)*

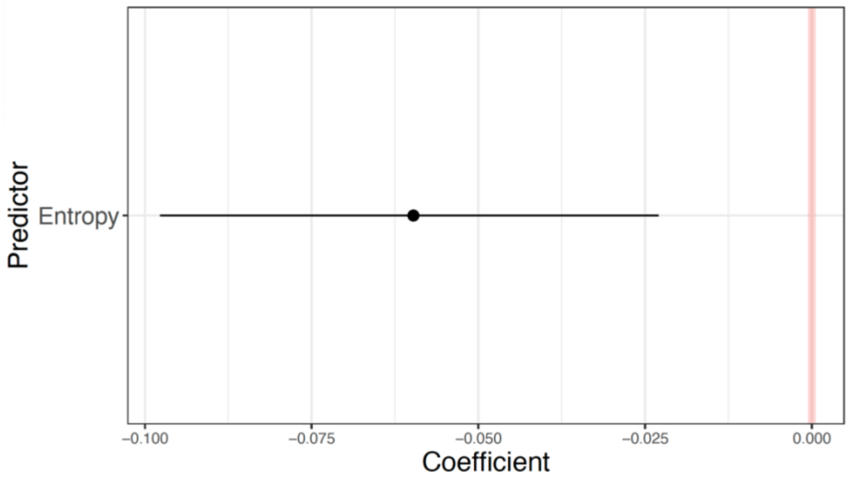


*Stimulation Dataset Winning Models (LOOIC and Pseudo-BMA methods agreed)*

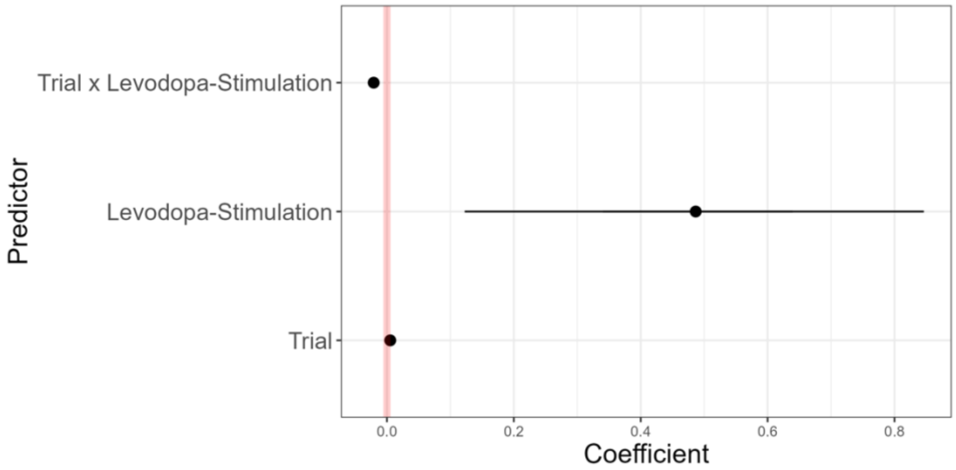


#### Table S5. Exploratory complete dataset selected model-coefficients

| Thought type | Active vs Sham Stimulation (Across Placebo Condition) | | Levodopa vs Placebo (Across Sham Condition) | | Active with Levodopa vs Active with Placebo | |
|---|---|---|---|---|---|---|
| | LOOIC | Pseudo-BMA | LOOIC | Pseudo-BMA | LOOIC | Pseudo-BMA |
| **Deliberately constrained thought** | Mod20 – 0.41<br>Mod19 – 0.18 | Mod19 – 0.74<br>Mod20 – 0.16 | Mod12 – 0.23<br>Mod19 – 0.21 | Mod19 – 0.91<br>Mod18 – 0.04 | Mod22 – 0.21<br>Mod19 – 0.16 | Mod19 – 0.78<br>Mod22 – 0.09 |
| **Automatically constrained thought** | Mod17 – 0.28<br>Mod21 – 0.19 | Mod21 – 0.26<br>Mod20 – 0.15 | Mod20 – 0.35<br>Mod17 – 0.18 | Mod20 – 0.37<br>Mod17 – 0.15 | Mod02 – 0.18<br>Mod21 – 0.15 | Mod18 – 0.46<br>Mod19 – 0.37 |

#### Table S6. Exploratory stimulation dataset selected model-coefficients

| Thought type | Active vs Sham Stimulation (Across Placebo Condition) | | Levodopa vs Placebo (Across Sham Condition) | | Active with Levodopa vs Active with Placebo | |
|---|---|---|---|---|---|---|
| | LOOIC | Pseudo-BMA | LOOIC | Pseudo-BMA | LOOIC | Pseudo-BMA |
| **Deliberately constrained thought** | Mod12 – 0.32 | Mod15 -0.40 | Mod13 – 0.34 | Mod13 – 0.75 | Mod15 – 0.54 | Mod13 – 0.33 |
| | Mod09 – 21 | Mod12 – 0.30 | Mod14 – 0.20 | Mod14 – 0.09 | Mod01 – 0.18 | Mod15 – 0.26 |
| **Automatically constrained thought** | Mod15 – 0.39 | Mod15 – 0.37 | | | Mod13 – 0.39 | Mod13 – 0.80 |
| | Mod11 – 0.35 | Mod14 – 0.23 | | | Mod10 – 0.28 | Mod10 – 0.04 |

#### Table S7. Winning LOOIC model from each set of exploratory probit models

A. Deliberately Constrained Thought – Active vs Sham (across Placebo conditions)

**Complete Dataset**

| Predictor | Estimate | 95% CIs |
|---|---|---|
| Behavioural variability | -.37 | [-.65, -.09]* |
| Randomness | -.01 | [-.06, .03] |
| Block | -.04 | [-.18, .09] |
| Trial | -.02 | [-.02, -.02]* |
| Behavioural variability x Randomness | -.20 | [-.39, -.01]* |

**Stimulation Block Dataset**

| Predictor | Estimate | 95% CIs |
|---|---|---|
| Trial | -.02 | [-.02, -.01]* |
| Stimulation | -.05 | [-.34, .25] |

B. Automatically Constrained Thought – Active vs Sham (across Placebo conditions)

**Complete Dataset**

| Predictor | Estimate | 95% CIs |
|---|---|---|
| Block | -.15 | [-.38, .07] |
| Trial | .01 | [.00, .01] |
| Stimulation | -.01 | [-.40, .37] |
| Block x Stimulation | -.01 | [-.32, .29] |

**Stimulation Block Dataset**

| Predictor | Estimate | 95% CIs |
|---|---|---|
| Behavioural variability | .01 | [-.29, .33] |
| Randomness | -.09 | [-.14, -.03]* |
| Trial | .01 | [.00, .01] |
| Stimulation | -.04 | [-.39, .34] |
| Behavioural variability x Randomness | .15 | [-.07, .37] |

C. Deliberately Constrained Thought – Levodopa vs Placebo (across Sham conditions)

**Complete Dataset**

| Predictor | Estimate | 95% CIs |
|---|---|---|
| Behavioural variability | -.34 | [-.56, -.11]* |
| Randomness | -.07 | [-.12, -.01]* |
| Block | -.26 | [-.46, -.06]* |
| Levodopa | .45 | [.04, .86]* |
| Behavioural variability x Randomness | -.17 | [-.34, -.01]* |
| Block x Levodopa | -.13 | [-.42, .16] |
| Randomness x Levodopa | .07 | [.00, .15]* |

**Stimulation Block Dataset**

| Predictor | Estimate | 95% CIs |
|---|---|---|
| Trial | -.02 | [-.03, -.01]* |
| Levodopa | .06 | [-.39, .56] |
| Trial x Levodopa | .01 | [.01, .02]* |

D. Automatically Constrained Thought – Levodopa vs Placebo (across Sham conditions)

**Complete Dataset**

| Predictor | Estimate | 95% Credible Intervals (CIs) |
|---|---|---|
| Behavioural variability | .00 | [-.24, .23] |
| Randomness | -.06 | [-.10, -.01]* |
| Block | -.13 | [-.29, .03] |
| Trial | .01 | [.00, .01]* |
| Behavioural variability x Randomness | .11 | [-.06, .29] |

E. Deliberately Constrained Thought – Active Stimulation with Levodopa vs with Placebo

**Complete Dataset**

| Predictor | Estimate | 95% CIs |
|---|---|---|
| Behavioural variability | -.23 | [-.43, -.03]* |
| Randomness | .04 | [.00, .08]* |
| Block | -.07 | [-.26, .12] |
| Trial | -.02 | [-.02, -.01]* |
| Behavioural variability x Randomness | .04 | [-.12, .20] |
| Levodopa | -.10 | [-.45, .25] |
| Block x Levodopa | -.17 | [-.43, .09] |

**Stimulation Block Dataset**

| Predictor | Estimate | 95% CIs |
|---|---|---|
| Behavioural variability | -.20 | [-.41, -.01]* |
| Randomness | .01 | [-.04, .05] |
| Trial | -.01 | [-.02, -.01]* |
| Levodopa | -.26 | [-.61, .08] |
| Behavioural variability x Randomness | .06 | [-.11, .23] |

F. Automatically Constrained Thought – Active Stimulation with Levodopa vs with Placebo

**Complete Dataset**

| Predictor | Estimate | 95% CIs |
|---|---|---|
| Randomness | -.06 | [-.10, -.02]* |

**Stimulation Block Dataset**

| Predictor | Estimate | 95% CIs |
|---|---|---|
| Trial | .01 | [.00, .01]* |
| Levodopa | .49 | [.05, .93]* |
| Trial x Levodopa | -.02 | [-.03, -.01]* |

#### Table S8. Exploratory Bayesian independent samples t-tests

| Thought type | Active vs Sham Stimulation (Across Placebo Condition) | Levodopa vs Placebo (Across Sham Condition) | Active with Levodopa vs Active with Placebo |
|---|---|---|---|
| | $BF_{01}$ | $BF_{01}$ | $BF_{01}$ |
| **Deliberately constrained thought** | 4.89 | 1.51 | 1.88 |
| **Automatically constrained thought** | 5.04 | 3.76 | 2.23 |
